# Recovering historical fish eDNA from museum-preserved Antarctic filter feeders via non-destructive metabarcoding

**DOI:** 10.1101/2025.06.07.656140

**Authors:** Gert-Jan Jeunen, Sadie Mills, Marc Bailie, Quentin Mauvisseau, Miles Lamare, Stefano Mariani, William Pearman, Monika Zavodna, Jackson Treece, Sara Ferreira, Neil J. Gemmell

**Author notes:** Corresponding author: Gert-Jan Jeunen School of Biological and Marine Sciences University of Plymouth Devon PL4 8AA United Kingdom.

## Abstract

1. Recent technical advances have significantly enhanced the value of museum specimens for molecular research, with metagenomic and metabarcoding approaches expanding further the utility of museum collections. However, given the finite number of specimens, there is a critical need to move past destructive DNA extraction approaches and to explore non-destructive techniques.
2. In this study, we evaluated the feasibility of extracting historical fish eDNA from the ethanol preservative used to store Antarctic museum specimens. We compared a variety of extraction methods – centrifugation, evaporation, filtration, and precipitation – using ten replicate samples per treatment for statistical analyses. To assess potential differences in preservative-derived eDNA recovery across different filter-feeding taxonomic groups, we included a Bryozoa and two Porifera, i.e., Demospongiae and Hexactinellida.
3. Comparative analyses with tissue biopsies revealed that the total genomic DNA was significantly reduced for non-destructive approaches. However, 10 ml ethanol filtration performed equal to, or in some instances, outperformed tissue biopsies when investigating the historical eDNA of Antarctic fish using a 16S rRNA metabarcoding approach, both for the number of species detected (α-diversity) and community characterisation (β-diversity).
4. Our findings demonstrate the potential of ethanol preservative as a valuable, non-destructive source of historical fish eDNA from museum-stored filter-feeding specimens. These results highlight the viability of non-destructive sampling for molecular research on museum collections, preserving specimen integrity while enabling biodiversity assessments. Further refinement of non-destructive eDNA extraction could expand its applicability across taxa, collection types, and preservation methods, ensuring the long-term sustainability of museum-based genomic research.

## 1 INTRODUCTION

Covering over 70% of the Earth’s surface, marine biomes are critical for sustaining life by supporting key ecosystem services, including oxygen production, carbon cycling, and for the provision of food and energy (Hoegh-Guldberg & Bruno, 2010). These essential services are increasingly at risk due to mounting anthropogenic pressures and climate change, which are accelerating species extinctions and driving shifts in community composition, ultimately disrupting ecosystem functioning (Lotze, 2021). To prevent further ecosystem degradation and facilitate ecosystem restoration, the implementation of effective marine biodiversity conservation is essential (Danovaro et al., 2021).

Understanding and managing marine ecosystems, particularly in the context of conservation and restoration, requires interpreting current biodiversity trends in light of accurate historical baselines and long-term ecological changes (Finnegan et al., 2015; Harnik et al., 2012; Lotze & Worm, 2009). In terrestrial ecosystems, extensive historical data have deepened our understanding of the impacts of both direct (Roberts et al., 2017) and indirect (Rick et al., 2013) human activities. In contrast, marine conservation has only recently begun to incorporate historical and ancient data sources, such as fossil records (Finnegan et al., 2015) and sediment cores (Finney et al., 2002), to reconstruct ecological baselines. However, these sources are often limited in availability (Willis et al., 2007) and are costly and logistically challenging to obtain (Kittinger et al., 2015). As a result, our understanding of historical anthropogenic impacts on marine ecosystems remains incomplete (Hoegh-Guldberg & Bruno, 2010; Kidwell, 2015; Norris et al., 2013). This gap is especially pronounced in remote regions like Antarctica, which, despite its isolation, has been increasingly affected over the past century by human activities such as fishing (Pinkerton & Bradford-Grieve, 2014), whaling (Aronson et al., 2011), and broader climate-driven changes (Parkinson, 2019).

Environmental DNA (eDNA) has emerged in the last decade (Ficetola et al., 2008) as a powerful method for ecosystem monitoring (Bowers et al., 2021; Takahashi et al., 2023), frequently surpassing traditional techniques in both sensitivity (Boussarie et al., 2018) and cost (Lugg et al., 2018). Aquatic eDNA surveys, however, are currently restricted to the exploration of contemporary biodiversity patterns due to high degradation rates of nucleic acids in marine environments (Collins et al., 2018). Hence, high temporal resolutions have been observed for eDNA surveys (Jensen et al., 2022). The exclusively contemporary nature of eDNA signals currently limits the establishment of historical ecological baselines through eDNA surveys, which are essential for improving marine ecosystem management and restoration (Harnik et al., 2012; Kowalewski et al., 2015; Norris et al., 2013).

Beyond water sampling, eDNA can also be obtained from filter feeders, such as marine sponges (Brodnicke et al., 2023; Harper et al., 2023; Jeunen, Lamare, et al., 2024; Mariani et al., 2019), sea anemones (Cunnington et al., 2024), bivalves (Weber et al., 2023), and crustaceans (Siegenthaler et al., 2019). These natural samplers accumulate eDNA from their surrounding biological community within their tissue matrix through filter feeding. Comparative studies with aquatic eDNA have demonstrated a high degree of similarity between both eDNA sources (Jeunen, Cane, et al., 2023; Jeunen, Lamare, et al., 2023), suggesting that natural samplers can provide spatial and temporal resolutions comparable to those of aquatic eDNA. While contemporary natural samples capture only eDNA from the present biological community (Cai et al., 2022; Harper et al., 2023), similar to aquatic eDNA, the extraction of historical eDNA from filter feeders stored in scientific collections offers a unique opportunity to reconstruct historical ecological baselines and detect temporal ecosystem shifts with unprecedented accuracy (Jeunen, Mills, et al., 2024).

Recently, historical eDNA was successfully retrieved from museum-stored sponges to reconstruct Antarctic biodiversity patterns through DNA extraction from tissue biopsies (Jeunen, Mills, et al., 2024). Hence, novel and improved molecular approaches, such as metabarcoding, expand further the utility of museum collections (Gold et al., 2024; Hahn et al., 2020; Nakahama, 2021; Raxworthy & Smith, 2021), from mere biological specimens to invaluable repositories of historical molecular data (Gold et al., 2024). Natural eDNA sampler specimens, in particular, represent an untapped resource, functioning as molecular time capsules that retain genetic signatures of past biological communities at the time and place of specimen collection (Jeunen, Mills, et al., 2024).

The growing demand for museum specimens in genomic research (Hahn et al., 2020; Holmes et al., 2016; McGaughran, 2020) highlights the need for non-destructive DNA extraction methods (Gold et al., 2024; Rohland et al., 2004; Shokralla et al., 2010). Traditional approaches often rely on tissue biopsies, which, while effective, permanently alter or destroy parts of the specimen (Hofreiter & Shapiro, 2012). Given that museum collections represent a finite and irreplaceable resource, minimising destructive sampling is essential (Hahn et al., 2020; Hofreiter & Shapiro, 2012; McGaughran, 2020). Moreover, aging effects, preservation-related degradation, and the risk of contamination further necessitate the development of optimised, minimally invasive protocols (Hahn et al., 2020). One promising alternative is the extraction of DNA from the preservation medium, such as the ethanol in which specimens are stored (Gold et al., 2024; Rohland et al., 2004; Shokralla et al., 2010). When collected from filter-feeding specimens, this approach could allow researchers to recover not only the genomic DNA of the specimen, but also historical eDNA without compromising specimen integrity.

In this study, we address the critical need for non-destructive DNA extraction methods to recover historical eDNA from museum-preserved filter-feeding specimens. We compared non-destructive approaches with traditional tissue-based methods, assessing their effectiveness in recovering total DNA concentration, as well as capturing biodiversity through historical eDNA metabarcoding. We incorporated a Bryozoa and two Porifera to assess potential differences in preservative-derived eDNA recovery across different filter-feeding taxonomic groups. Our findings provide new insights regarding the use of non-destructive DNA extraction on the preservative fluids in which valuable museum specimens are stored. Besides enhancing our capacity to investigate historical biodiversity and better understand the long-term impacts of human activities on marine ecosystems, non-destructive methods enable us to do so without depleting or damaging invaluable museum collections.

## 2 MATERIALS & METHODS

### 2.1 Experimental design

Non-destructive sampling of Antarctic filter feeders was performed by processing the ethanol in which the specimens were submerged, using four different techniques: centrifugation, evaporation, filtration (1 ml and 10 ml), and precipitation. Unless otherwise specified, 1 ml of ethanol was processed per replicate. To benchmark the effectiveness of non-destructive approaches for metabarcoding, comparisons were made with tissue biopsy samples prepared using two treatments: blotted to remove excess ethanol; and unblotted. Each treatment was replicated ten times per specimen to enable statistical comparisons.

To assess the suitability of non-destructive sampling across filter-feeding taxa, three museum specimens were selected. Two represented major Porifera classes with differing internal skeletal structures: one species of Demospongiae (spongin, sometimes with siliceous spicules) and one species of Hexactinellida (hexagonal silica spicules). The third specimen belonged to the phylum Bryozoa, a filter-feeding group not previously evaluated as a natural eDNA sampler (Figure 1; Table 1). All specimens were collected in 2008 from the Ross Sea, preserved in 99% ethanol when collected, and subsequently stored in 80% ethanol at the NIWA Invertebrate Collection (NIC).

**Figure 1:**
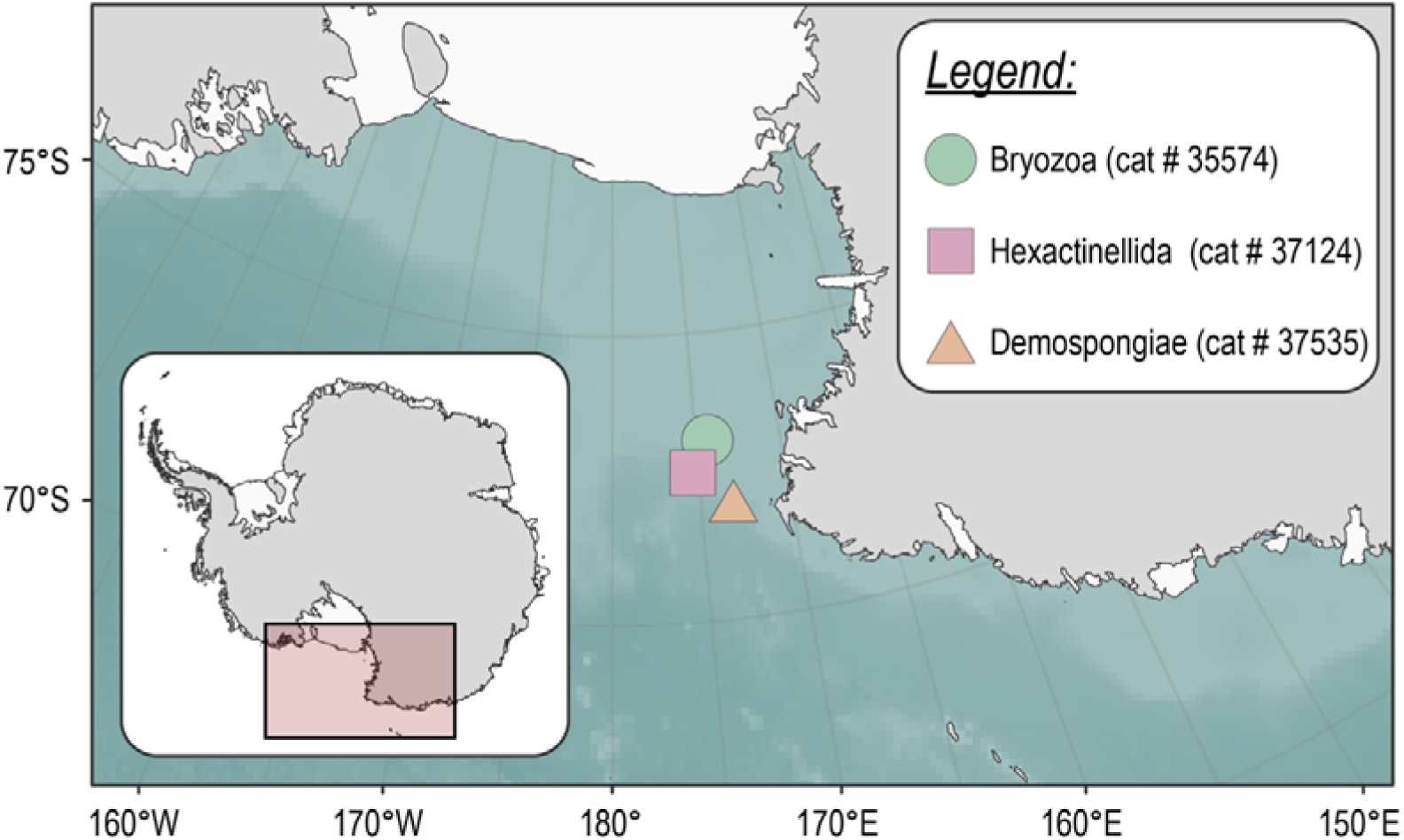
Map of the Ross Sea depicting collection points of museum specimens. Points are shaped and coloured by the taxonomic ID: Bryozoa (green circle), Hexactinellida (lilac square), and Demospongiae (beige triangle).

**Table 1:** Metadata of Antarctic museum specimens, including the NIWA Invertebrate Collection Catalogue number (CAT. #), phylum, class, and species identification based on morphology, the date, latitude, longitude, and depth.

| CAT. # | PHYLUM | CLASS | SPECIES | DATE | LAT. | LONG. | DEPTH |
| --- | --- | --- | --- | --- | --- | --- | --- |
| 35574 | Bryozoa | Stenolaemata | *Neofungella claviformis* | 09/02/2008 | -73.1245 | 174.3205 | 321 m |
| 37124 | Porifera | Hexactinellida | *Rossella fibulata* | 21/02/2008 | -72.5903 | 175.3423 | 479 m |
| 37535 | Porifera | Demospongiae | *Homaxinella* sp. | 24/02/2008 | -72.0093 | 173.2238 | 850 m |

### 2.2 Laboratory processing of Antarctic museum specimens

Prior to subsampling, specimen jars were inverted twice to distribute particulate matter evenly within the preservative medium. Twenty tissue biopsies and 140 ml ethanol were collected from each specimen at NIC. Subsamples were transported to the University of Otago’s PCR-free eDNA facilities at Portobello Marine Laboratory (PML) to minimise contamination risk during subsequent sample processing steps. Bench spaces and equipment were sterilised using a 10% bleach dilution (0.5% hypochlorite final concentration) and wiped with ultrapure water (UltraPure^TM^ DNase/RNase-Free Distilled Water, Invitrogen^TM^) before laboratory work (Prince & Andrus, 1992). Additionally, negative extraction controls were processed alongside samples during DNA extraction (50 µl ultrapure water) and negative no template controls were added during qPCR amplification (2 µl ultrapure water) to check for contamination during sample handling. DNA extractions for each treatment were performed using the Qiagen DNeasy Blood & Tissue Kit (Cat # 69506; Qiagen GmbH, Germany) following the manufacturer’s recommendations, with slight modifications for each treatment (Supplement 1). Total DNA concentration and purity were measured on Qubit Flex (Cat # Q32854; Qubit^TM^ dsDNA HS Assay Kit, ThermoFisher Scientific, US) and Denovix DS-11 FX+ (Cat # 30266629; Denovix, US).

Library preparation followed the protocol of (Jeunen, Mills, et al., 2024) to investigate Antarctic fish diversity using a ∼200 bp fragment of the 16S rRNA gene: Fish16SF: 5’-GACCCTATGGAGCTTTAGAC-3’ (Berry et al., 2017); Fish16S2R: 5’-CGCTGTTATCCCTADRGTAACT-3’ (Deagle et al., 2007). Briefly, a 10-fold dilution series (neat, 10-fold, 100-fold) was used to optimise input DNA for qPCR amplification and identify inhibitors and low-template samples (Murray et al., 2015). qPCR reactions consisted of a 20 µl volume, including 1× SensiFAST SYBR Lo-ROX MIX (Cat # BIO-94005; Meridian Bioscience, UK), 0.4 µmol/L of the forward and reverse primer (Integrated DNA Technologies, Australia), 2 µl of template DNA, and ultrapure water as required. Amplification was carried out in duplicate and the thermal profile included (i) an initial denaturation step of 95°C for 3 minutes, followed by (ii) 50 cycles of 30 seconds at 95°C, 30 seconds at 54°C and 45 seconds at 72°C, and terminated with (iii) a melt-curve analysis.

Libraries were prepared using a one-step amplification protocol using fusion primers (Berry et al., 2017). Fusion primers consisted of an Illumina adapter, a modified Illumina sequencing primer, a 6-8 bp barcode tag, and the template-specific forward or reverse primer. Amplification was conducted in duplicate and each sample was assigned a unique barcode combination with different forward and reverse barcodes in a single sample. The qPCR mastermix and thermal profile were unmodified from the dilution series test described above. Duplicate qPCR reactions were combined to reduce stochastic effects from PCR amplification (Alberdi et al., 2018). Samples were pooled into mini-pools based on end-point qPCR fluorescence, C_t_-values, and melt-curve analysis. Mini-pools were visualised using gel electrophoresis to confirm the presence of a single band, and the concentration was measured on Qubit. Equimolar pooling produced a single library for each Antarctic museum specimen. Due to differences in DNA concentration among samples and negative controls, the latter were spiked into the library to allow for optimal library concentration according to Illumina MiSeq® specifications. The final library was size selected using Pippin Prep (Cat # PIP0001; Sage Science, US and quantified using Qubit. All three libraries, one per Antarctic museum specimen containing 70 samples and 10 controls, were sequenced at the Otago Genomics Facility, University of Otago (New Zealand) on an Illumina MiSeq® instrument using MiSeq reagent kit v2 1×300 bp, with 5% PhiX spiked into the library to minimise issues associated with low-complexity libraries.

### 2.3 Bioinformatic and statistical analysis

Raw sequencing data consisted of a single fastq file per library. FastQC *v* 0.11.5 (Andrews, 2010) was used to check the quality of the raw sequencing files prior to bioinformatic processing. The bioinformatic pipeline encompassed demultiplexing and primer removal using cutadapt *v* 4.1 (Martin, 2011), while VSEARCH *v* 2.13.3 (Rognes et al., 2016) was used for quality filtering, dereplication, chimera removal, and denoising. A taxonomic ID was assigned to all Zero-radius Operational Taxonomic Units (ZOTUs) through a local blastn *v* 2.10.1+ search (Altschul et al., 1990) against a local curated reference database built using CRABS *v* 0.1.9 (Jeunen et al., 2022). Data was further curated after taxonomy assignment, including (i) artefact removal using *tombRaider v* 1.0 (Jeunen, Fernandes, et al., 2024), (ii) contamination filtering by applying a read abundance threshold based on the highest proportion of reads assigned to a ZOTU in a negative control sample, and (iii) removal of ZOTUs without a taxonomic ID or assigned to off-target taxonomic groups. Additionally, the need to rarefy sequence data was assessed through (i) differences in total read count between treatments using a one-way ANOVA, (ii) a positive correlation between total read count and number of observed ZOTUs using a generalized linear model (GLM) with a Poisson distribution and log link function, (iii) a lack of plateauing of rarefaction curves using the ampvis2 *v* 2.8.3 R package (Andersen et al., 2018), and (iv) curvature indices below the threshold value of 0.1 generated by the DivE *v* 1.3 R package (Laydon et al., 2014). Finally, species accumulation curves were drawn for three orders of Hill numbers (Alberdi & Gilbert, 2019) and sample coverage was calculated using the iNEXT.3D *v* 1.0.6 R package (Chao et al., 2021) to assess if sufficient sampling was achieved and equal sampling effort required normalisation.

Bayesian phylogenetic trees were created as described in Jeunen et al. (2024). Briefly, trees were generated using BEAST *v* 2.7.7 (Bouckaert et al., 2019) after aligning ZOTU sequences using DECIPHER *v* 2.28.0 (Wright, 2016). Phylogenetic tree construction was performed with a Markov chain Monte Carlo (MCMC) chain length of 10^8^ iterations, sampling trees every 10,000. Convergence of the MCMC chains and effective sample size was checked using TRACER *v* 1.7.2 (Rambaut et al., 2018). The maximum credibility tree from the posterior sample of phylogenetic time-trees with a burn-in percentage of 10% was identified through Tree Annotator *v* 2.7.6 (Bouckaert et al., 2019) and used for subsequent analyses.

All statistical analyses were performed in R *v* 4.3.1 and RStudio *v* 2023.06.1 + 524. Outliers from the Denovix DS-11 FX+ measurements were identified and removed using Grubbs test. Differences in DNA concentration and purity between extraction treatments for each museum specimen were evaluated using Welch’s ANOVA, followed by the Games-Howell *post hoc* test. Metabarcoding data were transformed to presence-absence prior to statistical analysis, given the lack of an established correlation between eDNA signal strength and species abundance for metabarcoding data (Fonseca, 2018). Faith’s Phylogenetic Diversity (PD) was calculated with the picante *v* 1.8.2 R package (Kembel et al., 2010) to investigate phylogenetic α-diversity. One-way ANOVA with Tukey’s HSD *post hoc* tests were used to determine significant differences between treatments within each museum specimen. To visualize taxon detection overlap between tissue and non-destructive DNA extraction treatments, Venn diagrams were generated with the eulerr *v* 7.0.2 R package (Larsson, 2024). Unweighted uniFrac distance matrices were computed for all samples and for subsets corresponding to individual museum specimens using the picante R package to investigate phylogenetic β-diversity. Principal Coordinates Analysis (PCoA) ordination was conducted using the vegan *v* 2.6-8 R package (Dixon, 2003). The ordination plot for all samples was simplified into a stratigraphic diagram using the primary PCoA axis, thereby enhancing readability compared to a 2D plot, with the riojaPlot *v* 0.1-20 R package (Juggins, 2013). For individual museum specimens, ordination plots were visualized as 2D graphs using the ggplot2 *v* 3.5.1 R package (Wickham, 2016). Significant differences in phylogenetic β-diversity were assessed with permutational ANOVA (PERMANOVA) followed by pairwise PERMANOVA, using the vegan and pairwiseAdonis *v* 0.4.1 (Martinez Arbizu, 2020) R packages, respectively. Multivariate homogeneity of group dispersion was tested using the vegan R package to assess variability in multivariate data across groups. To evaluate sample classification, a random forest multi-label classification model was applied using the community matrix as predictors. This analysis was conducted with the randomForest *v* 4.7-1.2 (Liaw & Wiener, 2002) and caret *v* 6.0-9.4 (Kuhn, 2008) R packages to classify samples by museum specimen (dataset: all samples; response variable: specimen ID) and by DNA extraction treatment (dataset: subset per museum specimen; response variable: DNA extraction treatment). A Classification and Regression Tree (CART) was generated with the party *v* 1.3-17 R package (Hothorn et al., 2006) to visualize the potential decision tree structure used by the random forest algorithm.

Raw sequencing data and metadata files are available on figshare (https://figshare.com/projects/MarsdenObjective1EthanolComparison/184573). All code with exact parameters and intermediary outputs used during bioinformatic processing, data curation, and statistical analysis are available on GitHub (https://github.com/gjeunen/marsden_obj1_non-destructive_heDNA).

## 3 RESULTS

### 3.1 Total genomic DNA concentration and purity

Total genomic DNA concentration differed significantly among treatments for all three museum specimens, according to Welch’s ANOVAs (Bryozoa: *F*_6,61_ = 58.612, *p* < 1e^-15^; Hexactinellida: *F*_6,59_ = 134.01, *p* < 1e^-15^; Demospongiae: *F*_6,58_ = 227.65, *p* < 1e^-15^). *Post hoc* Games-Howell tests revealed that, for each specimen, the total genomic DNA extracted from tissue samples was significantly higher compared to the DNA extracted from the ethanol in which the specimen was stored (Figure 2a – c). This difference ranged from 47x for Hexactinellida to 156x for Bryozoa and 335x for Demospongiae. Furthermore, significant differences in total DNA concentration were observed between ethanol treatments. However, the optimal non-destructive treatment, i.e., the method for which the highest total DNA concentration was measured, differed among specimens, with centrifugation and precipitation best for the bryozoan specimen, 10 ml filtration best for the glass sponge, and all treatments except for 1 ml filtration best for the demosponge (Figure 2a – c). Finally, no significant difference in total DNA concentration was observed among extracting DNA from a tissue biopsy with or without blotting the tissue dry. Significant differences among treatments were also observed for DNA purity (Figure 2d – i), whereby DNA extractions from ethanol showed higher variability between replicates compared to tissue DNA extracts. This variability, however, could have been influenced by the low total genomic DNA concentrations measured in ethanol extractions, which approach the lower sensitivity threshold of Denovix DS-11 FX+.

**Figure 2:**
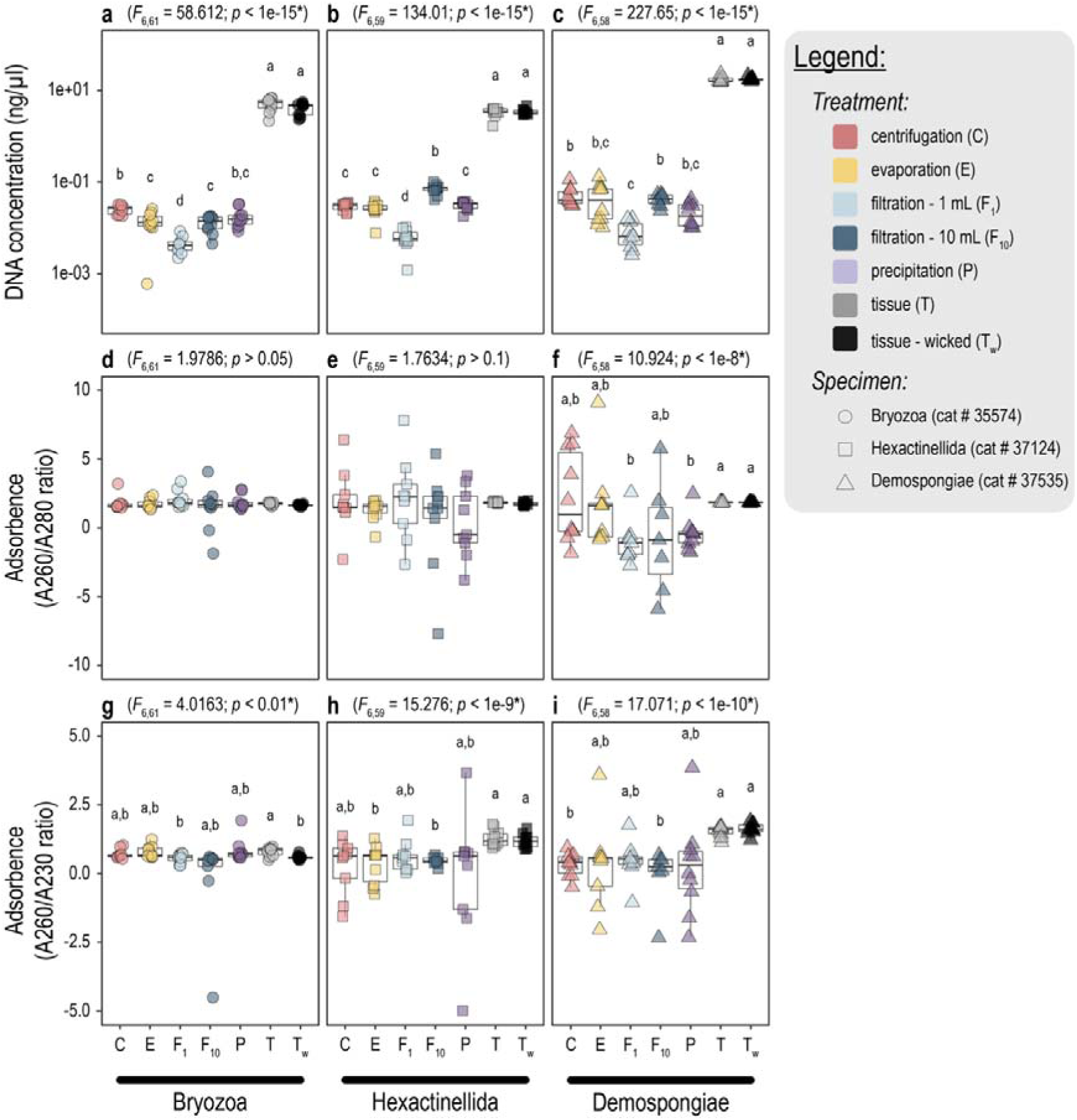
Boxplots depicting total DNA concentration (a – c), A260/A280 ratios (d – f), and A260/A230 ratios (g – i) for eDNA extracts from the bryozoan specimen (a, d, g), glass sponge specimen (b, e, h), and demosponge specimen (c, f, i). Measurements within boxplots are coloured based on extraction treatment, including centrifugation (C; red), evaporation (E; yellow), 1 ml filtration (F_1_; light blue), 10 ml filtration (F_10_; dark blue), precipitation (P; purple), tissue biopsy (T; grey), and blotted tissue biopsy (T_w_; black). The F-statistic and p-value for Welch’s ANOVA is provided above each facet, while post hoc Games-Howell test results are indicated by a letter above each boxplot whereby differing letters signify statistically significant differences between groups.

### 3.2 Metabarcoding sequencing results

Bioinformatic processing and data curation returned a total of 19,775,747 reads and 43 ZOTUs across all samples (Supplement 2). Samples were on average assigned 106,896 ± 5,057 reads. Data was not rarefied prior to statistical analysis, based on (i) non-significant differences in total read count between treatments, (ii) non-significant positive correlation between total read count and number of detected ASVs, (iii) plateauing of rarefaction curves, and (iv) all curvature indices above the threshold value of 0.1 (Supplement 3). Twenty-five (11.9%) out of 210 samples did not contain reads post data curation and were removed from subsequent analyses. All 25 samples came from the bryozoan specimen (cat # 35574), with most replicates from the tissue biopsy (blotting) treatment (9 out of 10 replicates), followed by tissue biopsy (7 out of 10 replicates), 1 ml filtration (5 out of 10 replicates), precipitation (2 out of 10 replicates), and evaporation (2 out of 10 replicates) treatments.

A single no template control for the bryzoan specimen (cat # 35574) was observed to contain a total of 57 reads (proportion: 2.8 × 10^-4^) assigned to five ASV’s, including 13 reads for ASV.1 (proportion: 3.0 × 10^-4^; taxonomic ID: *Chaenodraco wilsoni*), 4 reads for ASV.6 (proportion: 4.0 × 10^-4^; taxonomic ID: *Cryodraco atkinsoni*), 14 reads for ASV.7 (proportion: 1.8 × 10^-3^; taxonomic ID: *Pleuragramma antarctica*), 22 reads for ASV.14 (proportion: 1.42 × 10^-2^; taxonomic ID: *Laemonema* sp.), and 4 reads for ASV.21 (proportion: 3.9 × 10^-3^; taxonomic ID: *Chionodraco myersi*). ASV.14 was removed from the data set prior to statistical analysis, as it was only detected in a single sample besides the no template control. Furthermore, all detections not obtaining a proportion of 3.9 × 10^-3^ reads were set to 0 (based on the highest read abundance observed for the no template control, i.e., ASV.21), as well as singleton detections when the threshold was less than 1, to minimise the impact of background noise on statistical analyses. The number of samples collected for each treatment was deemed sufficient based on the flattening of species accumulation curves and sample coverage values ranging between 0.621 (treatment: 1 ml filtration; Bryozoa) and 0.979 (treatment: evaporation; Demospongiae) (Supplement 4).

### 3.3 Metabarcoding species detection

Our historical eDNA metabarcoding approach detected 43 unique taxa, including three Chondrichthyes and 40 Actinopterygii (Figure 3 a – c; Supplement 5). The three ASVs assigned to the class Chondrichthyes included two ASVs belonging to the genus *Bathyraja* within the order Rajiformes, as well as one member of the order Chimaeriformes. We also detected 12 families within the class Actinopterygii, including nine ASVs assigned to Nototheniidae, followed by seven ASVs assigned to Channichthyidae, six ASVs assigned to Myctophidae, and four ASVs assigned to Bathydraconidae. The most abundant taxon in our data, i.e., the ASV with the highest read count summed across all samples, was *Chaenodraco wilsoni* (ASV.1; 4,470,053 reads), followed by *Macrourus* sp. (ASV.2; 4,188,535 reads), *Trematomus* sp. (ASV.3; 2,789,541 reads), *Cryodraco antarcticus* (ASV.4; 2,562,992 reads), and *Bathylagus* sp. (ASV.5; 1,279,068 reads). The most frequently detected taxon, on the other hand, was *Pleuragramma antarctica* (ASV.20; number of detections: 100; proportion: 54.05%), followed by *Macrourus* sp. (ASV.2; number of detections: 82; proportion: 44.32%), *Trematomus* sp. (ASV.3; number of detections: 77; proportion: 41.62%), and *Cryodraco antarcticus* (ASV.4; number of detections: 74; proportion: 40.00%).

**Figure 3:**
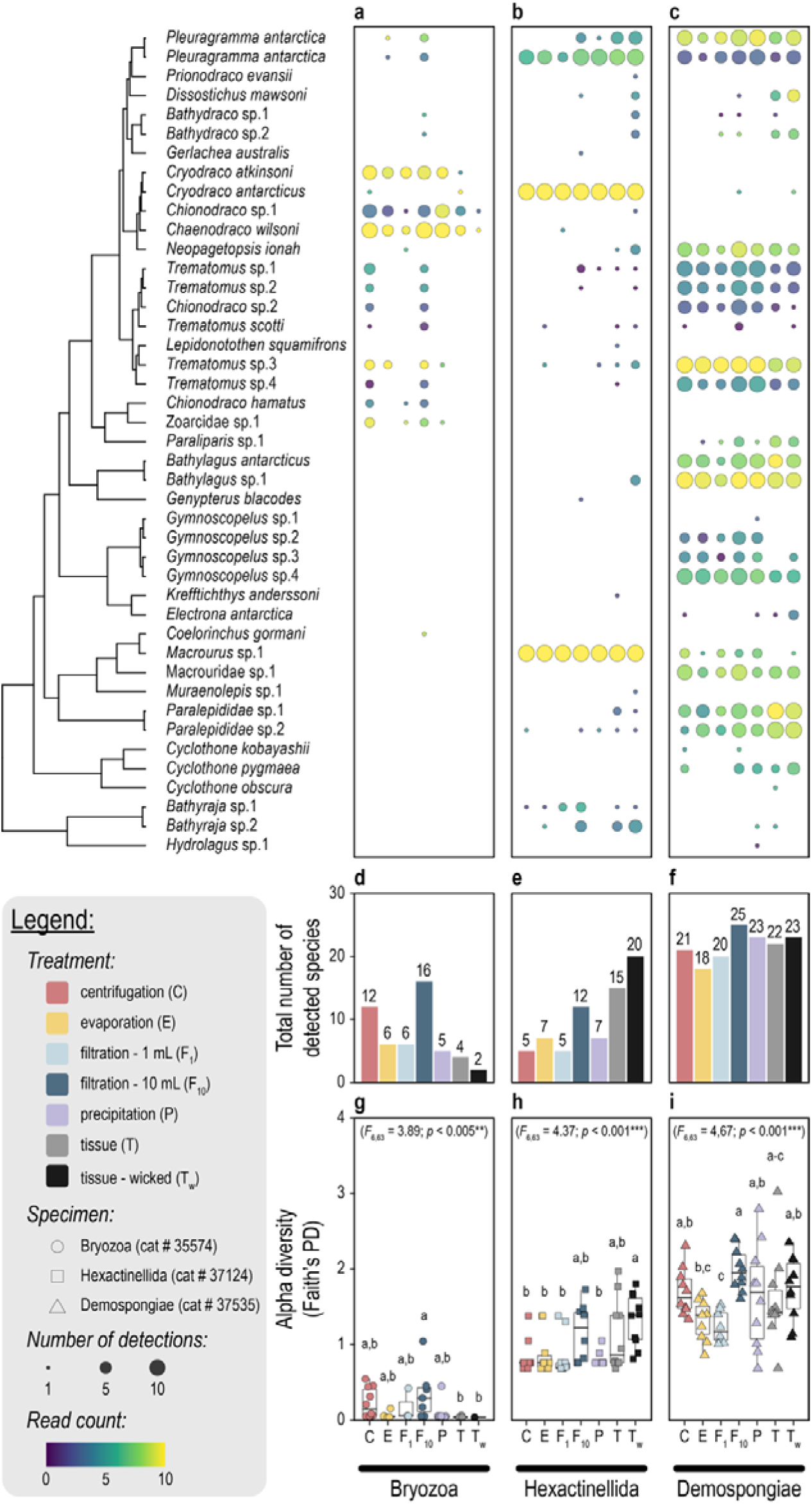
α-diversity analysis plots. Bayesian phylogenetic tree and bubble plots depicting number of detections by circle size and log-transformed read count by circle colour for the three specimens, including Bryozoa (a), Hexactinellida (b), and Demospongiae (c). Bar plots depicting total number of detected ASVs for each treatment per specimen (d – f). Number above bar represents the total number, while bars are coloured based on eDNA extraction treatment. Boxplots depicting Faith’s PD measurements for each treatment and specimen (g – i). One-way ANOVA results are presented at the top, while pairwise differences identified through post hoc Tukey HSD are indicated by a letter.

### 3.4 Phylogenetic alpha diversity

We detected varying numbers of species across specimens, with the demosponge yielding the highest diversity (30 species), followed by the glass sponge (26 species), and the bryozoan (18 species). Notably, no single treatment successfully detected all species identified across all treatments within any specimen. Among the non-destructive DNA extraction treatments, the filtration (10 ml) approach consistently outperformed the other treatments in terms of the total number of species detected across all ten replicates. Specifically, the filtration (10 ml) treatment detected 16 species (88.89%) in the bryozoan, 12 species (46.15%) in the glass sponge, and 25 species (83.33%) in the demosponge (Figure 3d – f). Furthermore, the filtration (10 ml) treatment outperformed both tissue-based extraction treatments in the bryozoan (filtration [10 ml]: 16; tissue: 4; tissue [wicked]: 2) and the demosponge (filtration [10 ml]: 25; tissue: 22; tissue [wicked]: 23). However, in the glass sponge, both tissue-based extractions detected more species overall compared to the filtration (10 ml) treatment (filtration [10 ml]: 12; tissue: 15; tissue [wicked]: 20).

Significant differences in Faith’s PD were observed between treatments within each specimen, as determined by one way ANOVA (Figure 3g – i). For the bryozoan (*F*_6,63_ = 3.886; *p* < 0.005**), the non-destructive 10 ml filtration DNA extraction treatment was observed to detect the largest number of ASVs per replicate on average (mean ± s.e.: 0.25 ± 0.10) and significantly outperformed both destructive tissue extraction treatments (tissue: 0.05 ± 0.01; tissue [wicked]: 0.04 ± 0.00) according to the *post hoc* Tukey HSD test. Although all other non-destructive DNA extraction treatments outperformed both tissue extraction treatments as well, the difference in Faith’s PD was found to be non-significant. For the glass sponge (*F*_6,63_ = 4.372; *p* < 0.001***), highest average Faith’s PD was observed for the tissue (wicked) extraction treatment (1.33 ± 0.11), with 10 ml filtration (1.16 ± 0.12) being the only non-destructive DNA extraction treatment that was found non-significantly different. For the demosponge (*F*_6,63_ = 4.674; *p* < 0.001***), the highest average Faith’s PD was observed for the non-destructive 10 ml filtration treatment (2.00 ± 0.10). While not significantly different from both destructive tissue treatments (tissue: 1.60 ± 0.19; tissue [wicked]: 1.74 ± 0.13), 10 ml filtration did perform significantly better than evaporation (1.32 ± 0.08) and 1 ml filtration (0.99 ± 0.17). Finally, no significant differences in Faith’s PD were observed between the two tissue-based extraction methods across all three specimens (Figure 3g – i).

### 3.5 Phylogenetic beta diversity

Non-destructive DNA extraction treatments recovered between 17.4% and 100% of ASVs identified by destructive tissue extraction methods within each specimen, as visualized by Venn diagrams (Supplement 6). For Bryozoa, centrifugation was the only treatment to recover all four taxa observed by destructive methods, due to the detection of *Cryodraco antarcticus*, while the other four non-destructive treatments identified three out of four taxa. In Hexactinellida, the filtration (10 ml) treatment recovered the most ASVs detected by destructive methods (10 ASVs; 43.5%), followed by precipitation (7 ASVs; 30.4%) and evaporation (7 ASVs; 30.4%). Notably, filtration (10 ml) and filtration (1 ml) were the only non-destructive treatments to detect ASVs not observed by destructive methods within the glass sponge, identifying two and one unique ASVs, respectively. For Demospongiae, filtration (10 ml) recovered the highest number of ASVs identified by destructive methods (23 ASVs; 88.5%), followed by precipitation (20 ASVs; 76.9%). Across all three specimens, filtration (10 ml) recovered the largest number of ASVs observed by destructive treatments (36 out of 53 ASVs; 67.9%), followed by precipitation (30 ASVs; 56.6%), centrifugation (28 ASVs; 52.8%), evaporation (27 ASVs; 50.9%), and 1 ml filtration (26 ASVs; 49.1%). Additionally, filtration (10 ml) also detected the highest number of ASVs not identified by destructive treatments (17 ASVs), followed by centrifugation (10 ASVs), 1 ml filtration (5 ASVs), precipitation (5 ASVs), and evaporation (4 ASVs).

Phylogenetic β-diversity differed significantly across all samples according to PERMANOVA for both the specimen (pseudo-*F*_2,184_ = 173.51; *p* < 0.001***) and DNA extraction (pseudo-*F*_6,184_ = 3.36; *p* < 0.001***) explanatory variables. Although both factors were highly significant, PERMANOVA revealed specimen ID (*R^2^* = 62.55%) as the largest explanatory variable for the observed variation in the dataset, while DNA extraction had a limited effect on community composition (*R^2^* = 3.64%). Significant differences in sample dispersion between specimens were also observed according to PERMDISP (*F*_2,184_ = 16.74; *p* < 0.001***), whereby pairwise comparisons revealed samples from the demosponge to have a higher average distance from the centroid compared to samples from the bryozoan and glass sponge specimens. PERMANOVA and PERMDISP results were corroborated by PCoA, which split samples originating from different specimens along the primary axis explaining 45.34% of the variation observed in the dataset (Figure 4a). Furthermore, the significant difference in phylogenetic β-diversity between specimens enabled the machine learning random forest classification model to successfully predict from which specimen a sample originated. The model achieved an overall out-of-bag (OOB) error rate of 1.08%, indicating high predictive accuracy. The confusion matrix revealed perfect classification for Bryozoa and Hexactinellida (class error = 0.00%), while Demospongiae exhibited a slightly higher error rate (class error = 2.86%) due to the misclassification of two samples as Bryozoa. Overall, the random forest model accurately predicted 183 of 185 samples (98.92% accuracy). The CART identified the presence of ASV.4 (*Cryodraco antarcticus*) as a significant indicator for the glass sponge, while the absence of ASV.4, but presence of ASV.1 (*Chaenodraco wilsoni*) as a significant indicator for the bryozoan and the absence of ASV.4 and ASV.1, but presence of ASV.5 (*Bathylagus* sp.) as a significant indicator for the demosponge (Supplement 7).

**Figure 4:**
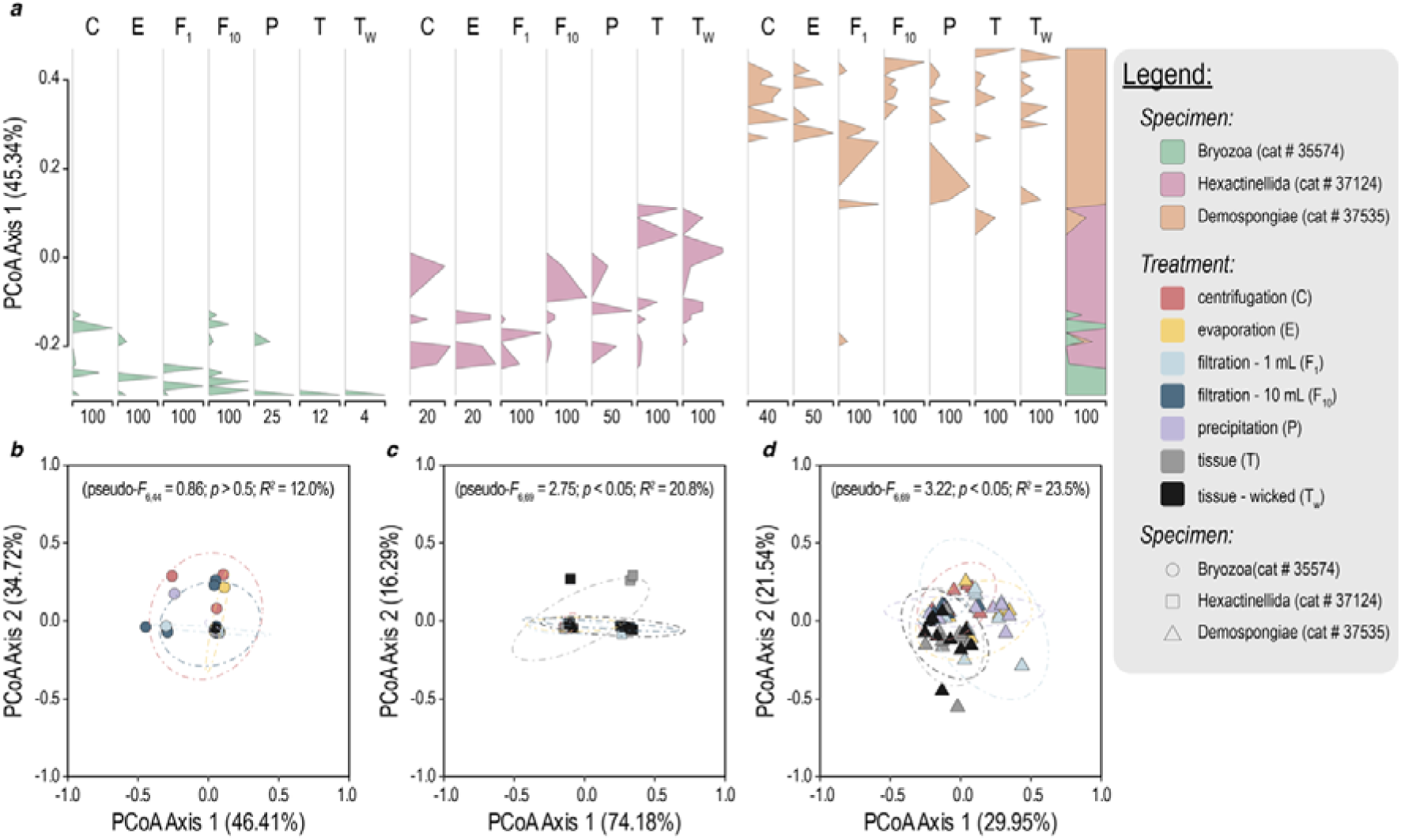
β-diversity analysis plots. (a) Stratigraphic diagram illustrating the primary PCoA axis for all samples. The x-axis represents the percentage of points distributed along the primary PCoA axis (y-axis). Facets represent DNA extraction treatments. The cumulative plot on the right-hand side aggregates the data across all facets. (b – d) PCoA plots for individual specimens: (b) Bryozoa, (c) Hexactinellida, and (d) Demospongiae. The percentage of variation explained by the primary and secondary PCoA axes are noted in the axis labels. Points are coloured by DNA extraction treatment and shaped by specimen. PERMANOVA results are reporting the pseudo-F statistic, p-value, and R^2^ value.

To eliminate the effect of specimen ID and more closely investigate differences between DNA extraction treatments, PERMANOVA, ordination, and random forest analyses were conducted for each specimen separately (Figure 4b – d). For all three specimens, PCoA revealed large sample overlap in 2D space without separating DNA extraction treatments along the primary (Bryozoa: 46.41%; Hexactinellida: 74.18%; Demospongiae: 29.95%) or secondary (Bryozoa: 34.72%; Hexactinellida: 16.29%; Demospongiae: 21.54%) axes. While PERMANOVA revealed no significant differences between DNA extraction treatments for the bryozoan specimen (pseudo-*F*_6,44_ = 0.86; *p* > 0.05; *R^2^* = 12.0%), significant differences were observed for the glass sponge (pseudo-*F*_6,69_ = 2.75; *p* < 0.01; *R^2^*= 20.8%) and demosponge specimens (pseudo-*F*_6,69_ = 3.22; *p* < 0.01; *R^2^* = 23.5%). Pairwise comparisons, however, revealed significant PERMANOVA differences to coincide with significant differences in sample variability, rather than placement in the 2D ordination space (Supplement 8). Finally, the lack of significant differences in phylogenetic β-diversity between DNA extraction treatments within each specimen limited the machine learning random forest classification model to predict from which DNA treatment a sample originated. The model achieved an OOB error rate of 75.56% for the bryozoan, 81.43% for the glass sponge, and 81.43% for the demosponge specimen (Supplement 9).

## 4 DISCUSSION

Our study pioneers an indirect, non-destructive approach to unlock historical biodiversity data from an overlooked resource, namely the ethanol preservative in which filter-feeding museum specimens are stored. While earlier work has shown the utility of investigating the composition of larvae fish assemblages from the ethanol these samples are stored in (Gold et al., 2024), our study goes a step further by reconstructing the biodiversity from the filtered eDNA by marine sponges and bryozoans which have been subsequently collected and stored in ethanol. By demonstrating that 10 ml filtration of ethanol recovers comparable Antarctic fish diversity to destructive tissue biopsies, scientists can employ powerful genomic techniques while safeguarding specimen integrity for future research as technologies advance (Hahn et al., 2020; Nakahama, 2021; Raxworthy & Smith, 2021). Our method has the potential to leverage the reported high spatiotemporal resolution of natural sampler eDNA (Brodnicke et al., 2023; Cai et al., 2022; Jeunen, Cane, et al., 2023; Mariani et al., 2019). This will unlock precise historical baselines and reveal undocumented ecosystem shifts with unprecedented accuracy, thereby providing critical data to guide conservation strategies in our rapidly changing oceans (Harnik et al., 2012; Lotze, 2021; Lotze & Worm, 2009). Hence, this approach is particularly valuable, not only for regions where baseline biodiversity data is scarce, but also by aligning with growing ethical and practical imperatives to minimise damage to irreplaceable specimens (Nakahama, 2021; Raxworthy & Smith, 2021).

The reduced total DNA concentration observed in ethanol extracts aligns with previous studies demonstrating the inherent trade-off between specimen preservation and DNA yield in non-destructive approaches (Rohland et al., 2004; Shokralla et al., 2010). However, our results challenge the assumption that higher total DNA quantities necessarily improve metabarcoding outcomes (Kawato et al., 2021; Ruan et al., 2022). Despite yielding significantly less DNA, ethanol-based extractions, particularly 10 ml filtration, showed no significant differences compared to tissue biopsies in detecting α- and β-diversity of Antarctic fish communities. This discrepancy likely reflects fundamental differences in DNA source partitioning. While tissue biopsies predominantly recover host genomic DNA, ethanol accumulates a mixture of leached host DNA and eDNA from the specimen’s historical environment. Hence, to better evaluate eDNA protocol optimisation studies, we recommend future research to incorporate targeted genomic measurements (e.g., qPCR assays for host vs. target taxa) alongside bulk DNA quantification (e.g., Qubit). This distinction is critical, not only for natural sampler and diet eDNA metabarcoding studies where host DNA often dominates (Pompanon et al., 2012), but also for conventional eDNA metabarcoding workflows, where non-target taxa (e.g., bacteria, phytoplankton) may contribute disproportionately to total DNA yields while providing little ecological insight into target communities (Stat et al., 2017). By decoupling DNA quantity from taxonomic relevance, scientists can optimise methods for ecological inference, rather than total DNA yield alone.

The largest difference in α- and β-diversity was observed among the three museum specimens, rather than the different extraction methods. For example, specimen ID accounted for 62.6% of the observed variation in fish community composition according to PERMANOVA, while CART and random forest successfully classified samples based on which specimen they originated from. Since the specimens in our study were collected at different locations, these results provide further evidence for the previously-reported high spatiotemporal resolution of natural sampler eDNA (Brodnicke et al., 2023; Neave et al., 2023). While our experimental design could not isolate species-specific effects from location variables, the ecological niches of detected fish species identified location to be the main driver of the observed differences. The deep-sea demosponge (collection depth: 850 m) harboured a distinct deep-sea fish assemblage dominated by deep-sea smelts (genus *Bathylagus*), mesopelagic lanternfishes (genus *Gymnoscopelus*), and bioluminescent bristlemouths (genus *Cyclothone*). In contrast, the bryozoan (collection depth: 321 m) and glass sponge specimens (collection depth: 479 m) collected on the Ross Sea continental shelf were characterised by icefishes (family Channichthyidae), which would be consistent with known biogeographic patterns in Antarctic fish distributions (Van de Putte et al., 2021). We recommend future studies to test co-occurring filter feeders from similar habitats to quantify taxonomic biases, a critical step for scaling museum-based reconstructions across collections.

Our comparison of four ethanol-based DNA extraction methods revealed significant variability in performance, underscoring the need for protocol optimisation akin to the extensive refinements achieved in aquatic eDNA (Deiner et al., 2015; Jeunen et al., 2018; Kumar et al., 2020; Turner et al., 2014) and, in lesser extent, for natural sampler eDNA (Harper et al., 2023). Notably, the optimal method varied among specimen ID when controlling for the volume processed. This suggests that standardising approaches may require taxon-specific adjustments, similar to how optimal aquatic eDNA protocols are influenced by the physical characteristics of the water to be filtered (Kumar et al., 2022). The most influential variable was processing volume, with 10 ml filtration outperforming all other treatments across all specimens for total number of species detected. This finding aligns with aquatic eDNA studies, where larger water volumes improve detection probabilities (Govindarajan et al., 2022; Hunter et al., 2019; Sepulveda et al., 2019). Our experimental design, however, only tested volume scalability for filtration due to the practical implementation advantages. While the other treatments could theoretically achieve similar gains with larger volumes, their implementation faces logistical constraints, e.g., high-capacity rotors for centrifugation and fume hoods for safe evaporation. Future investigations could pursue (i) quantitative thresholds for optimal ethanol volumes for each method, (ii) the incorporation of alternative methods to extract DNA from ethanol, and (iii) the development of high-throughput adaptations. These refinements would mirror the rigorous optimisation conducted for aquatic eDNA (Bowers et al., 2021) and, ultimately, enable robust temporal comparisons across historical specimens.

Beyond its non-destructive advantage, extracting historical eDNA from ethanol offers another significant practical benefit for biodiversity surveys. Our results demonstrate that ethanol-derived eDNA yields more consistent metabarcoding detections across all three specimens compared to tissue biopsies. Inter- and extrapolation analysis revealed this consistency could reduce replication effort by 42.7 – 62.9% to recover 90% of historical fish diversity (Supplement 4), directly translating to substantial cost savings in large-scale studies (Sanches & Schreier, 2020; Smart et al., 2016). This improved efficiency likely stems from ethanol’s capacity to homogenise eDNA through gentle mixing (Trimbos et al., 2021), effectively creating a uniform DNA suspension that circumvents the patchy distribution characteristic of natural sampler tissues (Jeunen, Mills, et al., 2024). Where natural sampler eDNA workflows require extensive replication to overcome intra-tissue heterogeneity (Jeunen, Mills, et al., 2024), ethanol’s mixing properties enable comparable community detection with fewer samples, reducing both processing time and sequencing expenses. For museum collections housing thousands of specimens, this methodological refinement could make comprehensive biodiversity assessments financially viable without compromising data quality.

While our results demonstrate that reduced DNA yields from ethanol do not impact metabarcoding success, reduced DNA yields could be a constraining factor for applications requiring high-input DNA, such as long-read sequencing technologies (Wang et al., 2021). Recent eDNA research has focused on implementing PacBio and Oxford Nanopore (Doorenspleet et al., 2025; Maggini et al., 2024; Stoeck et al., 2024) to sequence longer fragments for improved taxonomic resolution (Bylemans et al., 2018) and population-genetic analyses (Sigsgaard et al., 2020). The lower recovered DNA yield from ethanol extracts may limit this implementation of long-read sequencing beyond the expected shorter fragment length of historical specimens due to DNA degradation (Dabney et al., 2013). Future studies are required to evaluate whether ethanol-derived historical DNA could be successfully used with these novel technologies or if further DNA extraction optimisation is required to increase DNA yields from ethanol.

We propose future research to explore the broader applicability of non-destructive historical eDNA approaches across various taxa and preservative media. Our study contributes to this effort by encompassing both marine sponges, the most widely studied group for natural sampler eDNA (Cai et al., 2024; Gallego et al., 2024; Jeunen, Cane, et al., 2023; Mariani et al., 2019; Turon et al., 2020), and, for the first time, a bryozoan museum specimen. This novel inclusion of a bryozoan highlights the untapped potential of lesser-studied filter-feeding taxonomic groups as natural eDNA samplers. Preliminary work has already demonstrated that other filter feeders, including sea anemones (Cunnington et al., 2024), bivalves (Weber et al., 2023), and crustaceans (Siegenthaler et al., 2019), can also retain eDNA. By exploring the feasibility of extracting historical eDNA from different filter-feeding taxonomic groups, scientists could unlock a larger proportion of museum collections to reconstruct historical biodiversity baselines. Equally important is broadening the range of preservatives compatible with historical eDNA recovery. Although ethanol currently serves as a common preservation buffer in museum collections (Wandeler et al., 2007), recent advances in extraction chemistry now enable DNA retrieval from samples stored in e.g., formalin (Hahn et al., 2021). Adapting these protocols for historical eDNA analysis could incorporate valuable specimens before ethanol became standard (Wandeler et al., 2007). These advances would transform museum collections into global observatories of anthropogenic change, providing temporal baselines required to guide evidence-based conservation (Lotze & Worm, 2009).

## 5 CONCLUSION

Our study demonstrates that non-destructive DNA extraction from ethanol preservative, particularly 10 ml filtration, recovers Antarctic fish diversity comparable to destructive tissue biopsies while preserving museum specimens for future research. This approach addresses a critical challenge in genomic studies, i.e., balancing the growing demand for genetic data with the ethical imperative to conserve irreplaceable collections. We also show the potential for ethanol-based extractions to offer financial benefits over destructive methods, whereby increased detection consistency reduces replication requirements and processing costs, a significant advantage for large-scale biodiversity assessments.

By transforming museum collections into accessible repositories of historical eDNA, we establish filter-feeding specimens as powerful molecular time capsules. Our methodology facilitates reconstructions of past ecosystems without depleting finite collections, providing conservationists with two essential tools: (1) baselines to quantify anthropogenic impacts, and (2) temporal perspectives to distinguish natural variability from human-driven change. As we enter an era of accelerated biodiversity loss, integrating this approach into museum genomics workflows will expand the scientific and conservation value of collections while safeguarding them for future generations.

## 6 ACKNOWLEDGEMENTS

We thank the NIWA Invertebrate Collection (NIC) for providing access to the Antarctic specimens used in this study. We are grateful to Michelle Kelly, Jean Vacelet, and Dennis Gordon for the morphological identification of the specimens. We, also, thank the Otago Genomics Facility for the sequencing support provided throughout this study. This research was supported by funding from the Royal Society of New Zealand Marsden Fast-Start Fund (MFP-UOO2116), the Ministry of Business, Innovation, and Employment Antarctic Science Platform (MBIE ANTA1801), and a University of Otago Research Grant (UORG). We are grateful for the laboratory and technical staff at NIC and the University of Otago for their assistance during sample processing and data generation. We, also, thank the reviewers for their constructive feedback, which helped improve the clarity and quality of this manuscript. The authors declare no conflict of interest.

## 7 AUTHOR CONTRIBUTIONS

GJJ, SaM, ML, StM, and NJG designed the experiment. Funding was sourced by GJJ, SaM, and StM. Sample collection was performed by GJJ and SaM. Laboratory work was conducted by GJJ, MB, WP, MZ, JT, and SF. GJJ led the analysis, with help from SaM, MB, QM, ML, StM, WP, MZ, JT, SF, and NJG. GJJ wrote the manuscript, with significant input from SaM, QM, ML, StM, and NJG. All authors contributed to the writing of the manuscript.

## 8 DATA AVAILABILITY STATEMENT

Raw sequencing data and metadata files are available on figshare (https://figshare.com/projects/MarsdenObjective1EthanolComparison/184573). All code with exact parameters and intermediary outputs used during bioinformatic processing, data curation, and statistical analysis are available on GitHub (https://github.com/gjeunen/marsden_obj1_non-destructive_heDNA). The GitHub repository will be provided with a Zenodo DOI upon acceptance of this manuscript.

