## supplement for "Recovering historical fish eDNA from museum-preserved Antarctic filter feeders via non-destructive metabarcoding"

Supplement 1: DNA extraction protocols

All samples were extracted in the University of Otago’s PCR-free eDNA facilities at Portobello Marine Laboratory (PML) using the Qiagen DNeasy Blood & Tissue Kit. DNA extractions followed the manufacturer’s recommendations with slight modifications for each treatment. Below is a step-by-step guide of the DNA extraction protocol for each treatment, including (i) tissue biopsies, (ii) tissue biopsies with ethanol blotted away, (iii) filtration of 1 ml of ethanol, (iv) filtration of 10 ml of ethanol, (v) centrifugation of 1 ml of ethanol, (vi) precipitation of 1 ml of ethanol, and (vii) evaporation of 1 ml of ethanol. DNA extraction protocols did not differ between the three museum specimens and ten replicates were processed per treatment and specimen.

DNA extraction protocol: tissue biopsies

1. Cut 1 small piece of specimen (0.5 cm^3^) and place inside a 2 ml Eppendorf tube.
2. Add 720 µl Buffer ATL and 80 µl Proteinase K.
3. Vortex at maximum speed for 1 minute.
4. Incubate samples at 56°C overnight.
5. Remove samples from incubator and pipette mix 10 times.
6. Centrifuge at 6,000 x g for 1 minute at room temperature.
7. Pipette 600 µl supernatant into a new 2 ml Eppendorf tube.
8. Add 600 µl Buffer AL and vortex at maximum speed for 15 seconds.
9. Add 600 µl 100% ice-cold ethanol and vortex at maximum speed for 15 seconds.
10. Pipette the mixture into a DNeasy Mini Spin Column.
11. Centrifuge at 6,000 x g for 1 minute.
12. Discard the flow-through and collection tube.
13. Place the spin column in a new 2 ml collection tube.
14. Repeat steps 10 – 13 until all the mixture has gone through the spin column (change collection tubes every time and pipette maximum 620 µl of mixture each time).
15. Add 500 µl Buffer AW1 and centrifuge samples for 1 minute at 6,000 x g.
16. Discard the flow-through and collection tube.
17. Place the spin column in a new 2 ml collection tube.
18. Add 500 µl Buffer AW2 and centrifuge samples for 3 minutes at 20,000 x g.
19. Discard the flow-through and collection tube.
20. Place the spin column in a new 2 ml collection tube.
21. Centrifuge samples for 1 minute at 20,000 x g.
22. Discard the flow-through and collection tube.
23. Transfer the spin column in a 2 ml Eppendorf tube with caps removed.
24. Add 100 µl Buffer AE to the center of the membrane (preheat Buffer AE to 56°C).
25. Incubate samples for 1 minute at room temperature.
26. Centrifuge samples for 1 minute at 6,000 x g.
27. Repeat steps 24 – 26 for a total elution volume of 200 µl.
28. Transfer eluate to a new 1.5 ml Eppendorf tube.
29. Store DNA extracts at -20°C until further processing.

DNA extraction protocol: tissue biopsies with ethanol blotted away

1. Cut 1 small piece of specimen (0.5 cm^3^) and place inside a 2 ml Eppendorf tube.
2. Blot tissue biopsy on lint-free kim wipes to remove excess ethanol.
3. Continue with step 2 of protocol “DNA extraction protocol: tissue biopsies”.

DNA extraction protocol: 1 ml ethanol filtration

1. Invert specimen jar twice to distribute particulate matter evenly within ethanol.
2. Take a Wilderlab Standard eDNA Kit (filter pore size: 0.22 µm).
3. Take up 1 ml of ethanol using the Wilderlab 5 ml luer-lock syringe.
4. Press 1 ml of ethanol through the Wilderlab filter.
5. Add the DNA/RNA Shield buffer to the filter and store filter at 4°C for 1 week.
6. Remove buffer (~200 µl) from the filter and place in a new 2 ml Eppendorf tube.
7. Continue with step 2 of protocol “DNA extraction protocol: tissue biopsies”.

DNA extraction protocol: 10 ml ethanol filtration

1. Invert specimen jar twice to distribute particulate matter evenly within ethanol.
2. Take a Wilderlab Standard eDNA Kit (filter pore size: 0.22 µm)
3. Take up 10 ml of ethanol using the Wilderlab 50 ml luer-lock syringe.
4. Press 10 ml of ethanol through the Wilderlab filter.
5. Add the DNA/RNA Shield buffer to the filter and store filter at 4°C for 1 week.
6. Remove buffer (~200 µl) from the filter and place in a new 2 ml Eppendorf tube.
7. Continue with step 2 of protocol “DNA extraction protocol: tissue biopsies”.

DNA extraction protocol: 1 ml ethanol centrifugation

1. Invert specimen jar twice to distribute particulate matter evenly within ethanol.
2. Pipette 1 ml of ethanol into a 1.5 ml Eppendorf tube.
3. Centrifuge samples at 20,000 x g for 30 minutes.
4. Discard supernatant carefully. Ensure the pellet remains in the tube.
5. Add 180 µl of Buffer ATL and 20 µl of Proteinase K.
6. Vortex to mix sample thoroughly and incubate samples at 56°C overnight.
7. Vortex samples for 15 seconds, add 200 µl Buffer AL, and vortex for 15 seconds.
8. Add 200 µl of ice-cold ethanol (100%) and vortex sample for 15 seconds.
9. Continue with step 10 of protocol “DNA extraction protocol: tissue biopsies”.

DNA extraction protocol: 1 ml ethanol precipitation

1. Invert specimen jar twice to distribute particulate matter evenly within ethanol.
2. Pipette 1 ml of ethanol into a 1.5 ml Eppendorf tube.
3. Add 100 µl 5M NaCl to sample (14.6 g NaCl + PCR grade water to 50 ml volume).
4. Incubate at -20°C overnight.
5. Continue with step 3 of “DNA extraction protocol: 1 ml ethanol centrifugation”.

DNA extraction protocol: 1 ml ethanol evaporation

1. Invert specimen jar twice to distribute particulate matter evenly within ethanol.
2. Pipette 1 ml of ethanol into a 1.5 ml Eppendorf tube.
3. Open lid and place in dry-bath incubator at 56°C until fully evaporated.
4. Continue with step 5 of “DNA extraction protocol: 1 ml ethanol centrifugation”.

Supplement 2: metabarcoding read statistics

Illumina sequencing returned a total of 36,349,307 raw reads (Supplemental Figure 1), of which 24,149,058 (66.4%) reads could be assigned to samples during demultiplexing through cutadapt *v* 4.1 (Martin, 2011). Quality filtering using VSEARCH *v* 2.13.3 (Rognes et al., 2016) retained a total of 20,363,664 (56.0%) reads. Further denoising and chimera removal brought the total number of reads down to 20,247,244 (55.7%). The final number post data curation counted 19,775,747 (54.4%) reads. Similar demultiplexed read counts between samples (114,984.9 ± 74,363.1) and proportion of reads retained during various bioinformatic processing steps (quality filtering: 77.3% ± 28.7%; pre-data curation: 99.1% ± 2.0%; post-data curation: 87.6% ± 32.6%) indicate successful equimolar pooling of the library and data curation (Supplemental Figure 1A). Sample drop-out was only observed post data curation and was limited to 25 samples, all originating from the bryozoan specimen (Cat # 35574). Sample drop-out was mostly observed within the tissue biopsy (blotting) treatment (9 out of 10 replicates), followed by tissue biopsy (7 out of 10 replicates), 1 ml filtration (5 out of 10 replicates), precipitation (2 out of 10 replicates), and evaporation (2 out of 10 replicates).

A total of 2.231 (0.006%) reads were assigned to 15 negative control samples during demultiplexing (Supplemental Figure 1B). Negative control samples were assigned on average 148.7 ± 231.1 reads, with an extraction control assigned the highest number of reads of 730 (0.002%). Quality filtering removed 2,179 (97.7%) reads assigned to negative control samples, with only 57 (0.00028%) reads remaining within one no template control. These reads were assigned to five ZOTUs, including ZOTU 1 (taxon ID: *Chaenodraco wilsoni*; negative read count: 13; sample read count: 4,470,614; negative proportion: 0.0003%), ZOTU 6 (taxon ID: *Cryodraco atkinsoni*; negative read count: 4; sample read count: 985,377; negative proportion: 0.0004%), ZOTU 7 (taxon ID: *Pleuragramma antarctica*; negative read count: 14; sample read count: 786,855; negative proportion: 0.0018%), ZOTU 14 (taxon ID: *Laemonema* sp.; negative read count: 22; sample read count: 154,938; negative proportion: 0.0142%), and ZOTU 21 (taxon ID: *Chionodraco myersi*; negative read count: 4; sample read count: 102,528; negative proportion: 0.0039%). Given ZOTU 14 was only detected in a single sample besides the negative control, ZOTU 14 was removed from the entire data set prior to statistical analysis. The remainder of the data was filtered using an abundance threshold of 0.0039%, based on the highest read abundance observed for negative control samples, i.e., ZOTU 21.

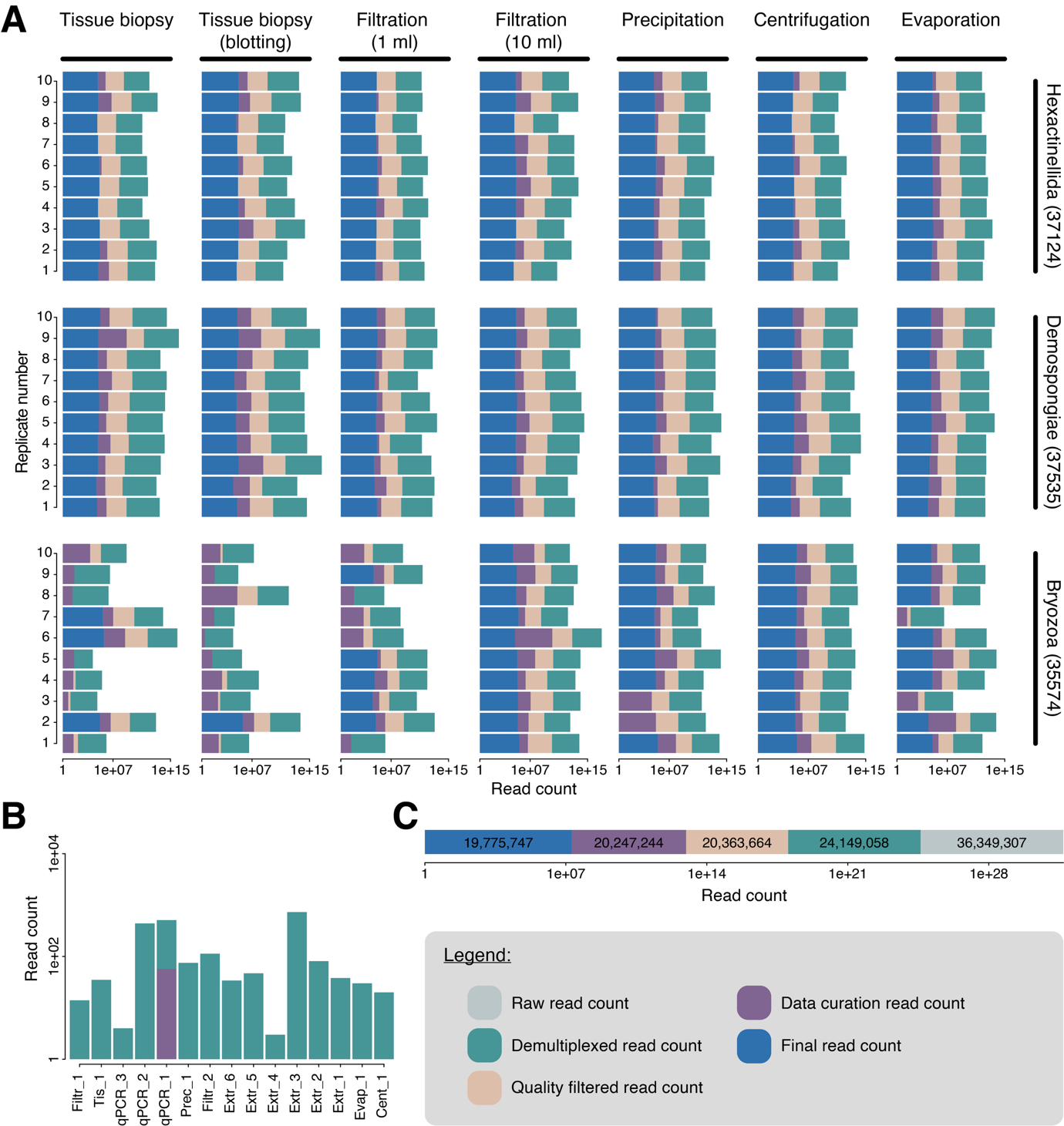

*Supplemental Figure 1: Stacked bar plots depicting read count during various stages of bioinformatic processing, including unfiltered (raw) reads (light green), demultiplexed read count (dark green), quality filtered read count (orange), pre-data curation read count (purple), and post-data curation read count (blue). The axis representing read count is log10 transformed for visualisation purposes. (A) Stacked bar plots depicting read count during various stages of bioinformatic processing for each sample separately, grouped by museum specimen and extraction treatment. X-axis depicts log10 transformed read count. Y-axis depicts replicate number. (B) Stacked bar plot depicting read count (log10-trasformed y-axis) during various stages of bioinformatic processing for negative control samples (x-axis). (C) Stacked bar plot depicting read count (log10-transformed x-axis) during various stages of bioinformatic processing for all samples and negative controls combined.*

Supplement 3: rarefaction analysis

Total read count was observed to be distributed evenly across samples and ranged between 21,576 reads and 532,384 reads for sample E35_1 (extraction treatment: Evaporation; specimen: Demospongiae; replicate: 1) and T74_6 (extraction treatment: Tissue biopsy; specimen: Bryozoa; replicate: 6), respectively (Supplemental Figure 2). Total read count across all samples averaged 106,896 ± 5,057. Total read count for samples collected from the bryozoan specimen received on average a higher read count (162,412 ± 13,497) compared to samples from the glass sponge (93,711 ± 2,984) and demosponge (84,392 ± 4,070). This difference in average read count between specimens was not significant, however, according to a one-way ANOVA (*F*_2,205_ = 1.312; *P* = 0.272).

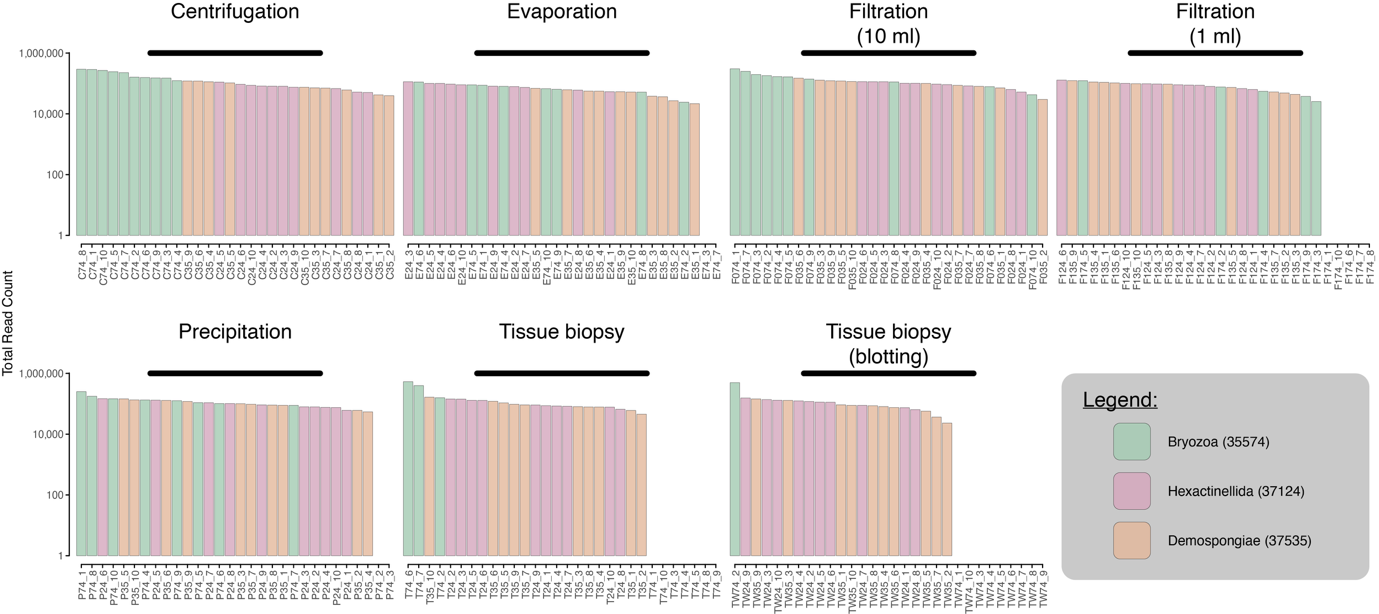

*Supplemental Figure 2: bar plots depicting total read count (log10-trasformed y-axis) post data curation per sample (x-axis). Bar plots are separated by extraction treatment and bars are coloured based on specimen ID, with samples from the bryozoan specimen in green, glass sponge in lilac, and demosponge in beige.*

A generalized linear model (GLM) was employed to examine the relationship between total read count ($y$) and number of detected ZOTUs ($z$). The model was specified with a Poisson distribution and log link function. The resulting equation can be expressed as:

$$\log\left( E\left[ z \right] \right)=\beta_{0}+ \beta_{1} \cdot y$$

where:

- $\beta_{0}=1.949 (intercept)$
- $\beta_{1}= -1.022 \times{10}^{-6} (slope for total read count)$

The model revealed a statistically significant negative association between total read count and number of detected ZOTUs, indicating that an increased read count for a sample would not necessarily be associated with a larger number of detected ZOTUs (Supplemental Figure 3). The null deviance of the model was 549.66 with 184 degrees of freedom, while the residual deviance was 544.69 with 183 degrees of freedom, yielding an Akaike Information Criterion (AIC) value of 1189.6.

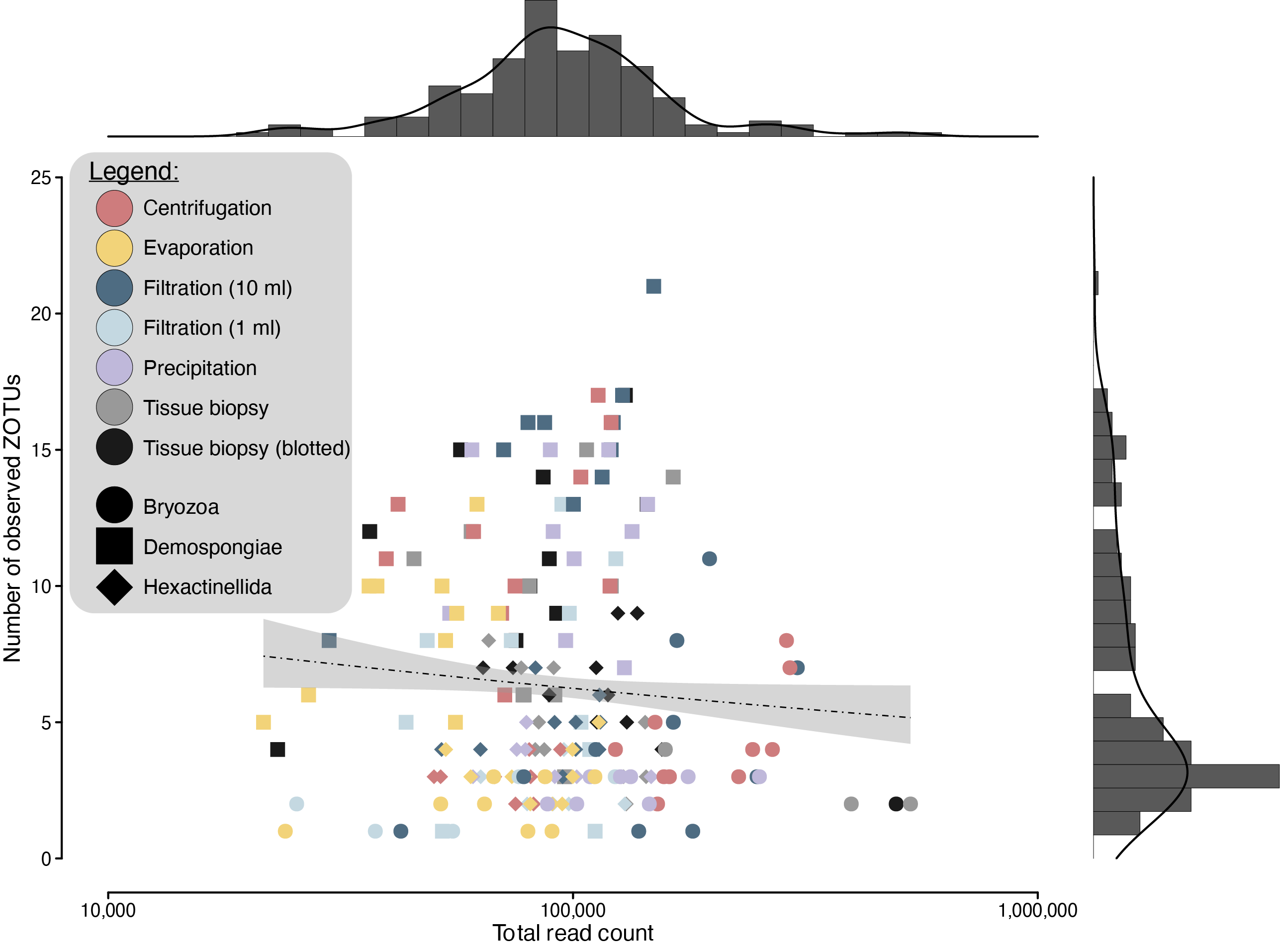

*Supplemental Figure 3: Scatter plot with total read count on the x-axis (log10 transformed) and number of detected ZOTUs on the y-axis. Shape is determined by specimen ID (Bryozoa: circle; Demospongiae: square; Hexactinellida: diamond), while colour is determined by extraction treatment (Centrifugation: red; Evaporation: yellow; Filtration [10 ml]: dark blue; Filtration [1 ml]: light blue; Precipitation: purple; Tissue biopsy: light grey; Tissue biopsy [blotted]: dark grey). The regression line is indicated by a dotted line and shaded area indicates the standard error to the regression line. Scatter plot is flanked by densigrams showing the distribution of sequencing depth and ZOTU richness across all samples.*

The plateauing of rarefaction curves indicates sufficient sequencing depth was achieved (Supplemental Figure 4). These results were corroborated by curvature indices, as no values were observed below the threshold of 0.1 (Supplemental Table 1).

*Supplemental Table 1: Curvature indices for each sample, with “REP” indicating the sample replicate. Curvature indices below the threshold of 0.1 are indicated by “*”. Samples dropped out during bioinformatic processing and data curation due to insufficient data are indicated by “NA”.*

| SPECIMEN | REP | TISSUE | TISSUE  (BLOT) | FILTER  (10 ML) | FILTER  (1 ML) | CENTR. | EVAP. | PRECIP. |
| --- | --- | --- | --- | --- | --- | --- | --- | --- |
| Bryozoa (35574) | 1 | NA | NA | 0.839 | NA | 0.878 | 0.963 | 0.944 |
|  | 2 | 0.993 | 0.956 | 1.000 | 0.994 | 0.972 | 1.000 | NA |
|  | 3 | NA | NA | 0.973 | 0.998 | 0.990 | NA | NA |
|  | 4 | NA | NA | 0.967 | 1.000 | 0.998 | 1.000 | 0.990 |
|  | 5 | NA | NA | 0.988 | 0.948 | 0.815 | 1.000 | 0.999 |
|  | 6 | 0.977 | NA | 0.971 | NA | 0.976 | 0.992 | 0.997 |
|  | 7 | 0.964 | NA | 0.995 | NA | 0.992 | NA | 0.798 |
|  | 8 | NA | NA | 0.998 | NA | 0.916 | 0.943 | 0.995 |
|  | 9 | NA | NA | 1.000 | 1.000 | 0.861 | 0.910 | 0.923 |
|  | 10 | NA | NA | 1.000 | NA | 0.995 | 0.928 | 0.858 |
| Hexactinellida (37124) | 1 | 0.998 | 0.946 | 0.980 | 0.999 | 0.996 | 0.988 | 0.999 |
|  | 2 | 0.999 | 0.993 | 0.995 | 0.999 | 0.999 | 0.998 | 0.999 |
|  | 3 | 0.947 | 0.984 | 0.997 | 0.996 | 0.986 | 0.991 | 0.902 |
|  | 4 | 0.992 | 0.871 | 0.910 | 0.999 | 0.999 | 0.999 | 0.999 |
|  | 5 | 1.000 | 0.990 | 0.876 | 0.999 | 0.967 | 0.999 | 0.999 |
|  | 6 | 0.999 | 0.926 | 0.998 | 1.000 | 0.998 | 0.999 | 0.999 |
|  | 7 | 0.937 | 0.989 | 0.836 | 0.997 | 0.997 | 0.999 | 0.999 |
|  | 8 | 0.790 | 0.993 | 0.995 | 0.999 | 0.996 | 0.998 | 0.997 |
|  | 9 | 0.988 | 0.999 | 0.998 | 0.999 | 0.999 | 0.999 | 0.999 |
|  | 10 | 0.990 | 0.982 | 0.999 | 0.999 | 0.999 | 0.999 | 0.997 |
| Demospongiae (37535) | 1 | 0.890 | 0.924 | 0.940 | 0.942 | 0.903 | 0.987 | 0.956 |
|  | 2 | 0.971 | 0.997 | 0.997 | 0.952 | 0.927 | 0.947 | 0.931 |
|  | 3 | 0.970 | 0.879 | 0.960 | 0.959 | 0.997 | 0.916 | 0.947 |
|  | 4 | 0.999 | 0.982 | 0.989 | 1.000 | 0.948 | 0.951 | 0.856 |
|  | 5 | 0.988 | 0.964 | 0.955 | 0.989 | 0.895 | 0.823 | 0.955 |
|  | 6 | 0.979 | 0.958 | 0.926 | 0.958 | 0.989 | 0.928 | 0.973 |
|  | 7 | 0.999 | 0.950 | 0.981 | 1.000 | 0.885 | 0.937 | 0.972 |
|  | 8 | 0.997 | 0.970 | 0.968 | 0.914 | 0.889 | 0.947 | 0.960 |
|  | 9 | 0.999 | 0.919 | 0.971 | 0.936 | 0.938 | 0.981 | 0.971 |
|  | 10 | 0.953 | 0.980 | 0.925 | 0.967 | 0.945 | 0.975 | 0.975 |

Sequencing data was not rarefied prior to statistical analysis based on (i) an even total read count distribution across samples (Supplemental Figure 2), (ii) non-significant positive correlation between total read count and number of detected ZOTUs (Supplemental Figure 3), (iii) plateauing of rarefaction curves (Supplemental Figure 4), and (iv) curvature indices were not observed below the threshold of 0.1 (Supplemental Table 1).

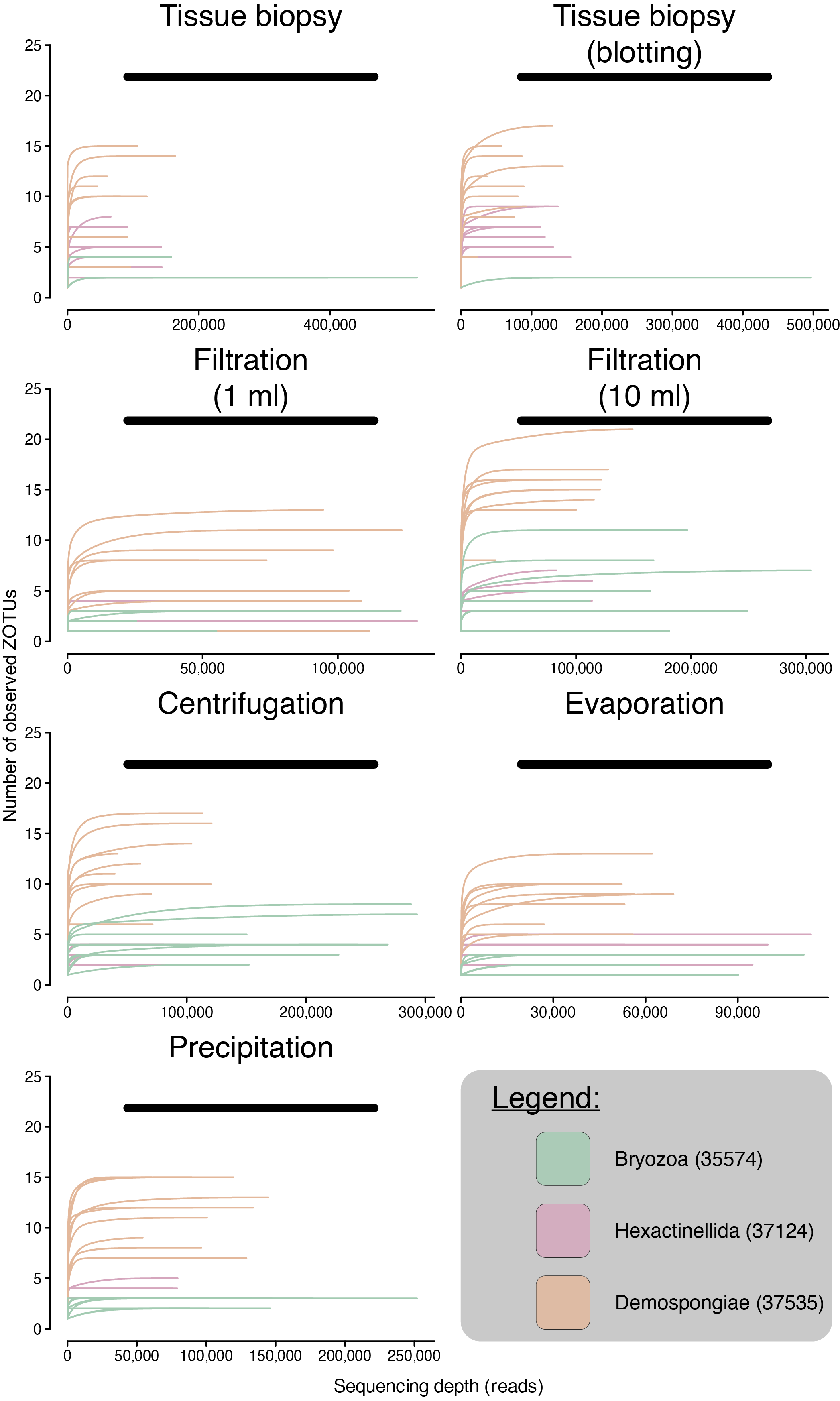

*Supplemental Figure 4: Rarefaction curves drawn for each sample, with number of reads on the x-axis and number of detected ZOTUs on the y-axis. Rarefaction curves are grouped by extraction treatment, i.e., tissue biopsy, tissue biopsy (blotted), filtration (1 ml), filtration (10 ml), centrifugation, evaporation, and precipitation. Rarefaction lines are coloured based on specimen ID, with the bryozoan specimen in green, demosponge in beige, and glass sponge in lilac.*

Supplement 4: species accumulation curves

Species accumulation curves were observed to flatten for all treatment groups across all three orders of Hill numbers (Supplemental Figure 5), indicating sufficient sampling was achieved within the experiment. Treatment groups consisted of the ten replicates per extraction method and museum specimen, while extrapolation enabled the investigation of double the number of samples within a treatment group, i.e., twenty samples.

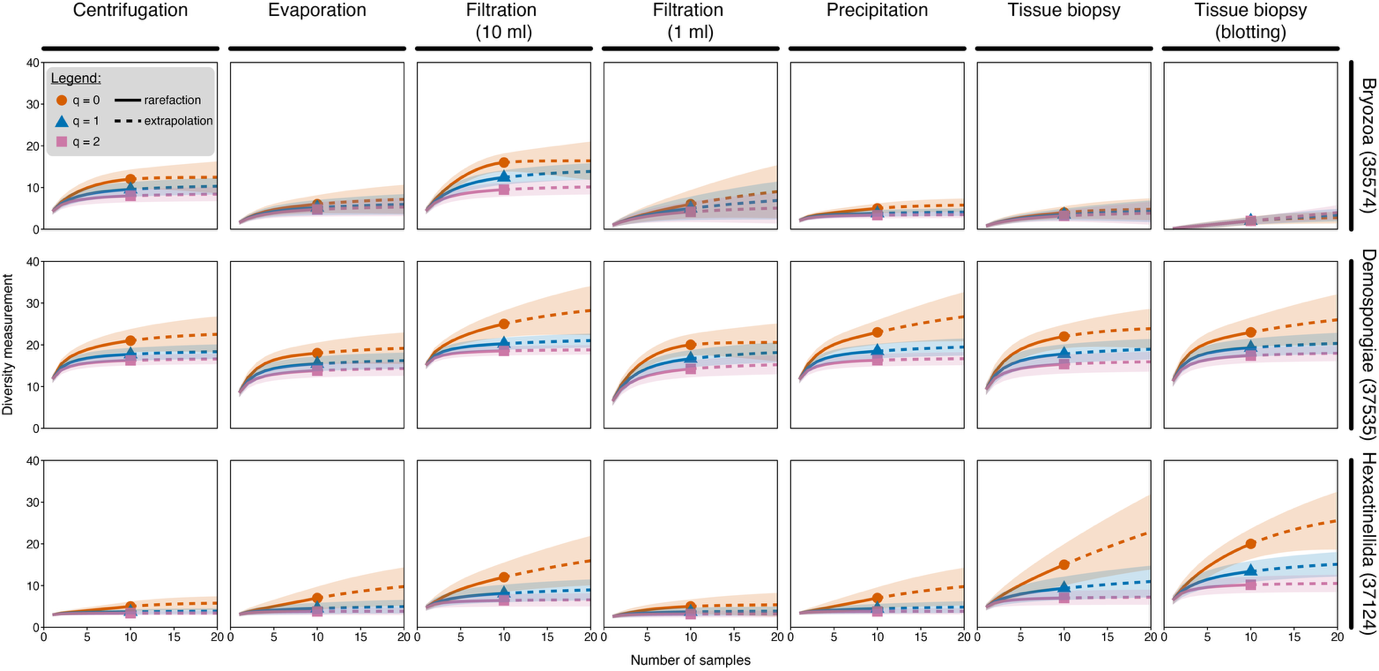

*Supplemental Figure 5: Species accumulation curves facetted by museum specimen (rows) and extraction treatment (columns) for three orders of Hill numbers, including q = 0 (species richness; orange), q = 1 (exponential of Shannon entropy; blue), and q = 2 (inverse of Simpson concentration; purple). Solid lines represent rarefaction analysis, while dotted lines represent extrapolation analysis up to double the number of samples collected.*

Further corroborating the findings from the species accumulation curves, was the high sample coverage observed per treatment group (Supplemental Table 2). Sample coverage ranged between 0.960 (precipitation treatment) and 0.979 (evaporation treatment) for the demosponge specimen, between 0.817 (tissue biopsy treatment) and 0.969 (1 ml filtration treatment) for the glass sponge specimen, and between 0.621 (1 ml filtration treatment) and 0.968 (centrifugation treatment) for the bryozoan specimen. Sample coverage could only not be calculated for the tissue biopsy (blotted) and tissue biopsy treatments of the bryozoan specimen, due to insufficient data from sample drop out.

*Supplemental Table 2: Sample Coverage for each group based on 10 replicates and a frequency-occurrence transformed data table format. Numbers represent percentage sample coverage between 0 and 1. NA values indicate insufficient data to calculate sample coverage, due to replicate drop out during bioinformatic processing and data curation.*

| TREATMENT | DEMOSPONGIAE | HEXACTINELLIDA | BRYOZOA |
| --- | --- | --- | --- |
| Tissue biopsy (blotted) | 0.966 | 0.874 | NA |
| Tissue biopsy | 0.963 | 0.817 | NA |
| Filtration (1 ml) | 0.970 | 0.969 | 0.621 |
| Filtration (10 ml) | 0.970 | 0.898 | 0.961 |
| Centrifugation | 0.978 | 0.945 | 0.968 |
| Precipitation | 0.960 | 0.890 | 0.922 |
| Evaporation | 0.979 | 0.880 | 0.888 |

Additionally, due to the generally higher sample coverage observed for ethanol over tissue treatments, less samples are estimated to be required to recover 90% of fish diversity within a specimen (Supplemental Table 3). The difference in number of required samples ranged between two (10 ml filtration treatment) and four (blotted tissue biopsy treatment) for the demosponge specimen, between five (1 ml filtration treatment) and twelve (blotted tissue biopsy treatment) for the glass sponge specimen, and between six (precipitation treatment) and fourteen (tissue biopsy treatment) for the bryozoan specimen.

*Supplemental Table 3: Estimated number of samples (± s.e.) required to recover 90% of diversity within the specimen x treatment. Extrapolation values are indicated by “*”.*

| TREATMENT | DEMOSPONGIAE | HEXACTINELLIDA | BRYOZOA |
| --- | --- | --- | --- |
| Tissue biopsy (blotted) | 3.647 (± 1.416) | 12.455 (± 6.832)* | 20.474 (± 1.033)* |
| Tissue biopsy | 4.971 (± 1.391) | 34.772 (± 12.912)* | 13.566 (± 1.664)* |
| Filtration (1 ml) | 6.975 (± 1.264) | 4.620 (± 0.865) | 34.640 (± 5.489)* |
| Filtration (10 ml) | 2.089 (± 1.078) | 10.423 (± 6.089)* | 8.370 (± 4.157) |
| Centrifugation | 2.768 (± 0.903) | 7.639 (± 0.432) | 6.720 (± 1.421) |
| Precipitation | 3.693 (± 1.514) | 11.274 (± 1.950)* | 5.633 (± 1.077) |
| Evaporation | 3.949 (± 0.937) | 12.567 (± 2.593)* | 11.118 (± 1.840)* |

Supplement 5: ASV table

The final ASV table post bioinformatic processing and data curation, including the taxonomic lineage and sequence for each ASV, on which all statistical analyses were conducted, can be found in Supplemental File “asv_table_final_with_tax_id_and_sequence.xlsx”.

Supplement 6: Venn diagrams

Venn diagrams were generated with the eulerr *v* 7.0.2 R package (Larsson, 2024) to visualize taxon detection overlap between tissue and non-destructive DNA extraction treatments (Supplemental Figure 6). A detection was set as “true” for a DNA extraction treatment if an ASV obtained a read count higher than 0 in at least one replicate sample. To simplify the analysis, replicates from both destructive DNA extraction treatments, i.e., tissue and tissue (wicked), were combined into a single treatment and compared against each of the five non-destructive DNA extraction treatments.

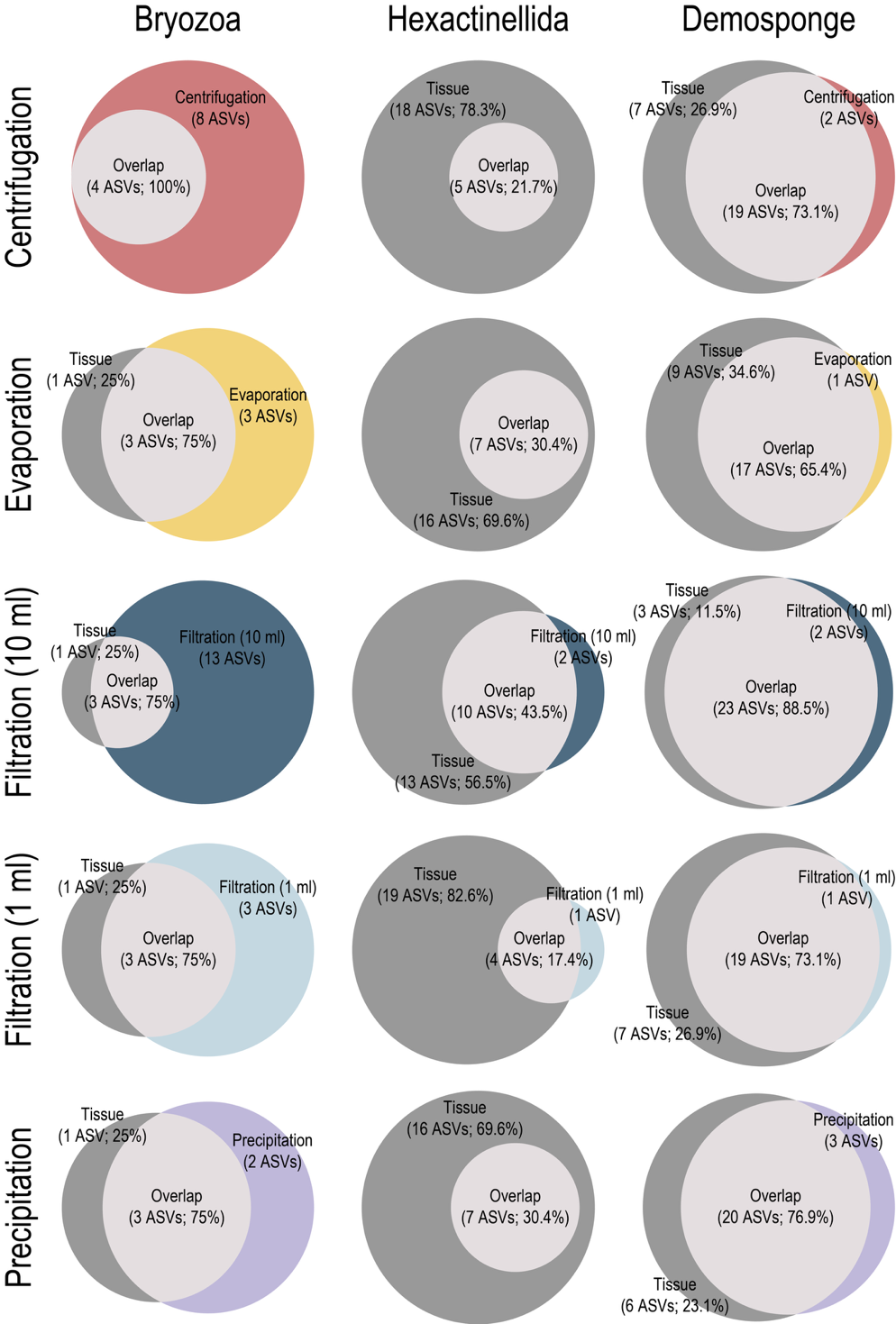

Supplemental Figure 6: Venn diagrams depicting ASV overlap between destructive and non-destructive DNA extraction treatments. Venn diagrams are grouped by specimen ID, with the bryozoan on the left, the glass sponge in the middle, and the demosponge on the right. Venn diagrams are grouped and coloured by non-destructive treatments, including centrifugation (first row; red), evaporation (second row; yellow), 10 ml filtration (third row; dark blue), 1 ml filtration (fourth row; light blue), and precipitation (fifth row; purple). The destructive treatment is coloured in dark grey, while the overlap between treatments is coloured in light grey. The size of Venn diagrams is proportional to ASV count.

Supplement 7: Classification and Regression Tree (CART)

The Classification and Regression Tree (CART), generated with the party *v* 1.3-17 R package (Hothorn et al., 2006), identifies the presence of ASV.4 (*Cryodraco antarcticus*) as indicative of samples originating from the glass sponge, while the absence of ASV.4 but presence of ASV.1 (*Chaenodraco wilsoni*) as indicative of samples originating from the bryozoan specimen, and the absence of ASV.4 and ASV.1 but presence of ASV.5 (*Bathylagus* sp.) as indicative of samples originating from the demosponge specimen (Supplemental Figure 7).

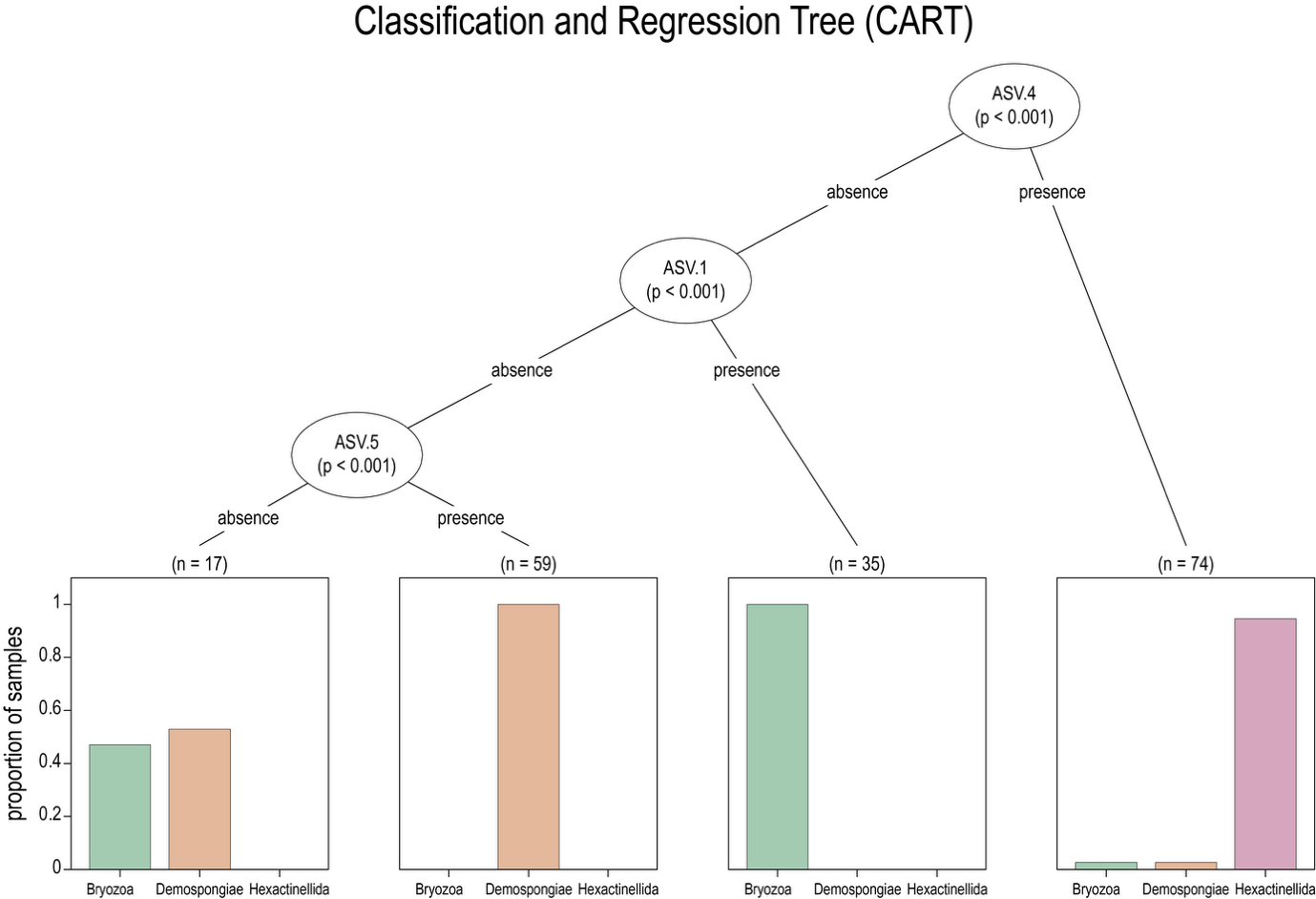

Supplemental Figure 7: CART based on the full metabarcoding dataset. P-values for each split, obtained through permutation, is provided in between brackets. “n” refers to the number of samples adhering to the decision. Bars are coloured based on specimen ID: Bryozoa (green), Hexactinellida (lilac), and Demospongiae (beige).

Supplement 8: Pairwise PERMANOVA and PERMDISP

Pairwise PERMANOVA and PERMDISP analyses were conducted with the pairwiseAdonis *v* 0.4.1 (Martinez Arbizu, 2020) and vegan *v* 2.6-8 (Dixon, 2003) R packages, respectively, to investigate differences in β-diversity between DNA extraction treatments within each museum specimen, including the bryozoan specimen (Supplemental Table 4), glass sponge (Supplemental Table 5), and demosponge (Supplemental Table 6).

Supplemental Table 4: Observed p-values for pairwise PERMANOVA (above diagonal) and PERMDISP (below diagonal) for each treatment within the bryozoan specimen, including centrifugation (C), evaporation (E), 1 ml filtration (F_1_), 10 ml filtration (F_10_), precipitation (P), tissue (T) and tissue wicked (T_W_). Significant p-values are highlighted in green.

| Bryozoa (cat # 35574) | | | | | | | |
| --- | --- | --- | --- | --- | --- | --- | --- |
|  | C | E | F_1_ | F_10_ | P | T | T_W_ |
| C |  | 0.096 | 0.353 | 0.659 | 0.316 | 0.263 | NA |
| E | 0.316 |  | 0.309 | 0.190 | 0.252 | 0.283 | NA |
| F_1_ | 0.323 | 0.844 |  | 0.541 | 0.511 | 0.578 | NA |
| F_10_ | 0.941 | 0.391 | 0.393 |  | 0.762 | 0.622 | NA |
| P | 0.073 | 0.364 | 0.581 | 0.109 |  | 0.716 | NA |
| T | 0.092 | 0.226 | 0.384 | 0.140 | 0.603 |  | NA |
| T_W_ | NA | NA | NA | NA | NA | NA |  |

Supplemental Table 5: Observed p-values for pairwise PERMANOVA (above diagonal) and PERMDISP (below diagonal) for each treatment within the glass sponge specimen, including centrifugation (C), evaporation (E), 1 ml filtration (F_1_), 10 ml filtration (F_10_), precipitation (P), tissue (T) and tissue wicked (T_W_). Significant p-values are highlighted in green.

| Hexactinellida (cat # 37124) | | | | | | | |
| --- | --- | --- | --- | --- | --- | --- | --- |
|  | C | E | F_1_ | F_10_ | P | T | T_W_ |
| C |  | 0.777 | 0.246 | 0.053 | 0.763 | 0.121 | 0.011 |
| E | 0.483 |  | 0.591 | 0.072 | 0.375 | 0.179 | 0.021 |
| F_1_ | 0.755 | 0.743 |  | 0.067 | 0.001 | 0.172 | 0.019 |
| F_10_ | 0.005 | 0.055 | 0.036 |  | 0.032 | 0.948 | 0.512 |
| P | 0.425 | 0.132 | 0.310 | <0.001 |  | 0.071 | 0.002 |
| T | 0.096 | 0.272 | 0.189 | 0.850 | 0.022 |  | 0.364 |
| T_W_ | 0.014 | 0.075 | 0.050 | 0.691 | <0.001 | 0.685 |  |

Supplemental Table 6: Observed p-values for pairwise PERMANOVA (above diagonal) and PERMDISP (below diagonal) for each treatment within the demosponge specimen, including centrifugation (C), evaporation (E), 1 ml filtration (F_1_), 10 ml filtration (F_10_), precipitation (P), tissue (T) and tissue wicked (T_W_). Significant p-values are highlighted in green.

| Demospongiae (cat # 37535) | | | | | | | |
| --- | --- | --- | --- | --- | --- | --- | --- |
|  | C | E | F_1_ | F_10_ | P | T | T_W_ |
| C |  | 0.012 | 0.011 | 0.259 | 0.308 | 0.002 | 0.004 |
| E | 0.957 |  | 0.144 | 0.002 | 0.074 | 0.002 | 0.001 |
| F_1_ | 0.004 | 0.004 |  | 0.001 | 0.14 | 0.002 | 0.001 |
| F_10_ | 0.252 | 0.229 | <0.001 |  | 0.04 | 0.009 | 0.043 |
| P | 0.026 | 0.028 | 0.256 | 0.003 |  | 0.002 | 0.015 |
| T | 0.021 | 0.022 | 0.484 | 0.003 | 0.706 |  | 0.723 |
| T_W_ | 0.039 | 0.042 | 0.152 | 0.005 | 0.754 | 0.508 |  |

Supplement 9: Random forest confusion matrices

A random forest multi-label classification model was applied using the community matrix as predictors and DNA extraction treatments as response for each museum specimen separately to investigate the impact of DNA extraction treatments on eDNA metabarcoding results. Analyses were conducted using the randomForest *v* 4.7-1.2 (Liaw & Wiener, 2002) and caret *v* 6.0-9.4 (Kuhn, 2008) R packages. Output printed within RStudio displaying the OOB and confusion matrix is provided below for each specimen.

Bryozoan specimen:

randomForest(formula = method ~ ., data = count_meta_bryozoa, mtry = 10, ntree = 10000)

- Type of random forest: classification
- Number of trees: 10000
- No. of variables tried at each split: 10
  - OOB estimate of error rate: 75.56%
- Confusion matrix:

|  | C | E | F_10_ | F_1_ | P | T | T_W_ | Class.error |
| --- | --- | --- | --- | --- | --- | --- | --- | --- |
| C | 1 | 1 | 3 | 1 | 3 | 1 | 0 | 0.900 |
| E | 0 | 2 | 0 | 2 | 4 | 0 | 0 | 0.750 |
| F_10_ | 4 | 0 | 4 | 0 | 2 | 0 | 0 | 0.600 |
| F_1_ | 1 | 2 | 0 | 1 | 1 | 0 | 0 | 0.800 |
| P | 4 | 0 | 1 | 0 | 3 | 0 | 0 | 0.625 |
| T | 1 | 0 | 0 | 0 | 2 | 0 | 0 | 1.000 |
| T_W_ | 0 | 0 | 0 | 0 | 1 | 0 | 0 | 1.000 |

Glass sponge specimen:

randomForest(formula = method ~ ., data = count_meta_hexactinellida, mtry = 14, ntree = 10000)

- Type of random forest: classification
- Number of trees: 10000
- No. of variables tried at each split: 14
  - OOB estimate of error rate: 81.43%
- Confusion matrix:

|  | C | E | F_10_ | F_1_ | P | T | T_W_ | Class.error |
| --- | --- | --- | --- | --- | --- | --- | --- | --- |
| C | 0 | 1 | 0 | 2 | 7 | 0 | 0 | 1.0 |
| E | 1 | 0 | 1 | 3 | 4 | 0 | 1 | 1.0 |
| F_10_ | 2 | 0 | 2 | 0 | 2 | 3 | 1 | 0.8 |
| F_1_ | 1 | 0 | 0 | 7 | 2 | 0 | 0 | 0.3 |
| P | 8 | 0 | 1 | 0 | 1 | 0 | 0 | 0.9 |
| T | 0 | 0 | 4 | 1 | 3 | 0 | 2 | 1.0 |
| T_W_ | 0 | 2 | 2 | 0 | 1 | 2 | 3 | 0.7 |

Demosponge specimen:

randomForest(formula = method ~ ., data = count_meta_demosponge, mtry = 16, ntree = 10000)

- Type of random forest: classification
- Number of trees: 10000
- No. of variables tried at each split: 16
  - OOB estimate of error rate: 81.43%
- Confusion matrix:

|  | C | E | F_10_ | F_1_ | P | T | T_W_ | Class.error |
| --- | --- | --- | --- | --- | --- | --- | --- | --- |
| C | 2 | 1 | 3 | 1 | 2 | 1 | 0 | 0.8 |
| E | 1 | 1 | 0 | 3 | 1 | 4 | 0 | 0.9 |
| F_10_ | 2 | 0 | 3 | 0 | 1 | 2 | 2 | 0.7 |
| F_1_ | 1 | 1 | 1 | 3 | 2 | 0 | 2 | 0.7 |
| P | 2 | 1 | 4 | 1 | 1 | 1 | 0 | 0.9 |
| T | 0 | 4 | 1 | 0 | 1 | 1 | 3 | 0.9 |
| T_W_ | 0 | 1 | 2 | 1 | 0 | 4 | 2 | 0.8 |
